# Prox2+ and Runx3+ neurons regulate esophageal motility

**DOI:** 10.1101/2022.07.07.499111

**Authors:** Elijah D. Lowenstein, Pierre-Louis Ruffault, Aristotelis Misios, Kate L. Osman, Huimin Li, Rebecca Thompson, Kun Song, Stephan Dietrich, Xun Li, Nikita Vladimirov, Jean-François Brunet, Andrew Woehler, Niccolò Zampieri, Ralf Kühn, Shiqi Jia, Gary R. Lewin, Nikolaus Rajewsky, Teresa E. Lever, Carmen Birchmeier

**Affiliations:** Developmental Biology/Signal Transduction, Max Delbrück Center for Molecular Medicine, Berlin, Germany and NeuroCure Cluster of Excellence, Charité-Universitätsmedizin Berlin, Corporate Member of Freie Universität Berlin and Humboldt-Universität zu Berlin, Berlin, Germany; Systems Biology of Gene Regulatory Elements, Berlin Institute for Medical Systems Biology, Max Delbrück Center for Molecular Medicine, Berlin, Germany; Department of Otolaryngology - Head & Neck Surgery, University of Missouri School of Medicine, Columbia, MO, USA; The First Affiliated Hospital, Jinan University, Guangzhou, China; Max Delbrück Center for Molecular Medicine, Berlin, Germany; Department of Cancer Research, Max Delbrück Center for Molecular Medicine, Berlin, Germany; Berlin Institute for Medical Systems Biology, Max Delbrück Center for Molecular Medicine, Berlin, Germany; Institut de Biologie de l’ENS (IBENS), Inserm, CNRS, École normale supérieure, PSL Research University, Paris, France; Genome Engineering & Disease Models, Max Delbrück Center for Molecular Medicine, Berlin, Germany; Department of Neuroscience, Max Delbrück Center for Molecular Medicine, Berlin, Germany and Neurowissenschaftliches Forschungszentrum, NeuroCure Cluster of Excellence, Charité-Universitätsmedizin Berlin, Corporate Member of Freie Universität Berlin and Humboldt-Universität zu Berlin, Berlin, Germany

**Author notes:** Lead Contact. Electronic address.

**Keywords:** Vagus nerve, vagal afferents, single cell RNA sequencing, esophagus, peristalsis, mechanoreceptors, mechanosensation, motility, homeostasis, digestion

## Abstract

Sensory neurons of the vagus nerve monitor distention and stretch in the gastrointestinal tract. Major efforts are underway to assign physiological functions to the many distinct subtypes of vagal sensory neurons. Here, we used genetically guided anatomical tracing, optogenetics and electrophysiology to identify and characterize three vagal sensory neuronal subtypes expressing Prox2 and Runx3. We show that these neuronal subtypes innervate the esophagus and stomach where they display regionalized innervation patterns. Their electrophysiological analysis showed that they are all low threshold mechanoreceptors, but possess different adaptation properties. Lastly, genetic ablation of Prox2+ and Runx3+ neurons demonstrated their essential roles for esophageal peristalsis and swallowing in freely behaving animals. Our work reveals the identity and function of the vagal neurons that provide mechanosensory feedback from the esophagus to the brain, and could lead to better understanding and treatment of esophageal motility disorders.

## Introduction

Sensory signaling through the vagus nerve relays vital information from internal organs to the brain and controls various aspects of bodily homeostasis (Brookes et al., 2013; Prescott and Liberles, 2022). During food intake and digestion, the ingested bolus exerts mechanical force against the walls of the digestive tract. Vagal sensory neurons located bilaterally in the nodose ganglia (also called the inferior vagal ganglia) sense stretch and tension in the digestive tract and transmit information on the location and size of the bolus to the brain (Mercado-Perez and Beyder, 2022). This mechanical information from the gut triggers the physiological and behavioral responses needed for food intake such as gut motility and appetite control (Kim et al., 2022).

Several types of mechanosensory neurons that display unique electrophysiological properties and form distinct types of endings like intraganglionic laminar endings (IGLEs), intramuscular arrays (IMAs) and mucosal endings have been detected in the gastrointestinal tract by both electrophysiological and histological techniques (Page et al., 2002; Phillips and Powley, 2000; Wang and Powley, 2000; Zagorodnyuk and Brookes, 2000; Zagorodnyuk et al., 2003). Due to the vast heterogeneity of vagal sensory neurons, their molecular characteristics were only recently defined by the use of single cell (sc) RNA-seq analyses (Bai et al., 2019; Kupari et al., 2019; Prescott et al., 2020). Further, work over the past few years has begun to characterize the properties of some subtypes of neurons important for digestion using molecular and genetic tools. For example, two subtypes of vagal neurons that form IGLEs contacting intestinal and stomach enteric ganglia express *Oxtr* and *Glp1r*, respectively. The activation of Oxtr+ intestinal IGLEs potently inhibits feeding, whereas Glp1r+ stomach IGLEs detect stretch (Bai *et al*., 2019; Williams et al., 2016). Further, recent work assigned Piezo2+Grm5+Slit2+ neurons as esophageal IMAs that respond to stretch (Zhao et al., 2022). IGLEs are also known to innervate esophageal enteric ganglia, but their molecular signature is currently unknown.

Swallowing (or deglutition) is an active process that transports food and liquid from the mouth to the stomach via the esophagus. The esophageal phase of swallowing requires peristaltic movements of the esophageal wall, which are controlled by reflexes that are executed by vagal motor neurons in the hindbrain (dorsal motor nucleus of the vagus and nucleus ambiguus), and the enteric ganglia in the esophagus (Ertekin and Aydogdu, 2003; Goyal and Chaudhury, 2008). Esophageal motility disorders are characterized by dysphagia, and are frequent comorbidities of aging and age-related neurological disease (Aslam and Vaezi, 2013; Aziz et al., 2016; Suttrup and Warnecke, 2016). The etiology underlying dysphagia is mostly unknown, due in large part to the lack of knowledge about the sensorimotor circuits involved (Kloepper et al., 2020). The first studies of esophageal physiology were undertaken in the late 19^th^ century, and for the following decades it was debated whether the activity of hindbrain swallowing centers (i.e. hindbrain vagal motor nuclei and the nucleus of the solitary tract) suffices to generate esophageal peristalsis, or if vagal sensory feedback is also required (Janssens et al., 1976; Jean, 1984; Kronecker H., 1883). Several lines of evidence subsequently demonstrated the importance of peripheral feedback for esophageal function, but the exact vagal subtype that provides this feedback is unknown (Falempin et al., 1986; Frazure et al., 2021; Lang, 2009). Classical studies often relied on blunt dissection of the vagus nerve and therefore affected both sensory and motor vagal fibers. Furthermore, the majority of these studies analyzed swallowing under anesthesia, which is known to affect esophageal peristalsis (Lang, 2009). Deglutition in awake, freely behaving rodents can be observed using videofluoroscopic swallowing studies (VFSS) (Haney et al., 2019; Hinkel et al., 2016; Lever et al., 2015). Thus, ablation of specific vagal sensory neurons that innervate the esophagus, and subsequent analysis using VFSS, can conclusively demonstrate the extent to which sensory feedback regulates esophageal peristalsis and define the responsible vagal sensory neuronal subtype.

Here, we used single cell RNA-sequencing (scRNA-seq) to characterize heterogeneity of vagal mechanosensory neurons, and identified subtype specific markers useful for genetic analysis. We found that the majority of Phox2b+ vagal neurons that express the mechanosensitive ion channel *Piezo2* are defined by their expression of two transcription factors, either *Prox2* or *Runx3*, and we provide evidence that neurons expressing these two factors are developmentally related. Cell-selective optogenetic activation, neuronal tracing and anatomical analyses were used to characterize the Prox2+/Runx3+ vagal neurons, which demonstrated that they innervate the esophagus and stomach forming IGLEs on enteric ganglia. The electrophysiological analysis demonstrated that Prox2+/Runx3+ neurons are highly sensitive to mechanical stimuli and are low-threshold mechanoreceptors. Lastly, we ablated these cells using intersectional genetic tools and performed videofluoroscopic swallowing analysis in awake and unrestrained animals, which revealed that these neurons are required for esophageal motility. Our data demonstrate the existence of specific vagal neuronal subtypes that control esophageal peristalsis and body homeostasis by relaying mechanosensory information from the esophagus to the brain.

## Results

### Molecular characterization of vagal mechanosensory neuronal types

To perform scRNA-seq, we isolated vagal sensory neurons using flow cytometry after genetically labeling all vagal neurons with a nuclear GFP reporter (ganglia from 15 *VGlut2^Cre^;R26^nGFP^* mice at postnatal day (P) 4, see Figure S1A). We used the CEL-Seq2 technology, a low throughput method with high gene detection sensitivity, and sequenced 1536 cells (Hashimshony et al., 2016; Kim et al., 2020b). After removing cells with more than 250,000 or less than 17,000 unique molecular identifiers, 1,392 cells remained for which a median of 9,323 genes and 70,667 unique molecular identifiers per cell were identified. The analysis of this dataset defined 22 distinct neuronal subtypes (Figure S1B; see Methods for further details). Our data were integrated with two scRNA-seq datasets of vagal neurons published during this study to perform a meta-analysis on 4,442 vagal neurons (Figure 1A, Figure S1B-D) that included 1392 neurons isolated at P4 (this study), 395 and 956 neurons isolated at P42 and between P56-84, respectively (Bai et al., 2019), and 1,707 neurons isolated between P32-35 (Kupari *et al*., 2019). Uniform Manifold Approximation and Projection (UMAP) analysis of the combined datasets revealed a large heterogeneity between vagal neurons and improved the resolution of their diversity. The meta-analysis indicated that the roughly 10,000 vagal neurons per mouse (Figure S1A) segregate into 27 transcriptomically distinct subtypes (Figure 1A, Figure S1C). Cells from all datasets contributed to all 27 subtypes, indicating that shortly after birth all subtypes are specified (Figure S1D). The meta-analysis can be freely explored at https://shiny.mdc-berlin.de/VGIE/.

**Figure 1.**
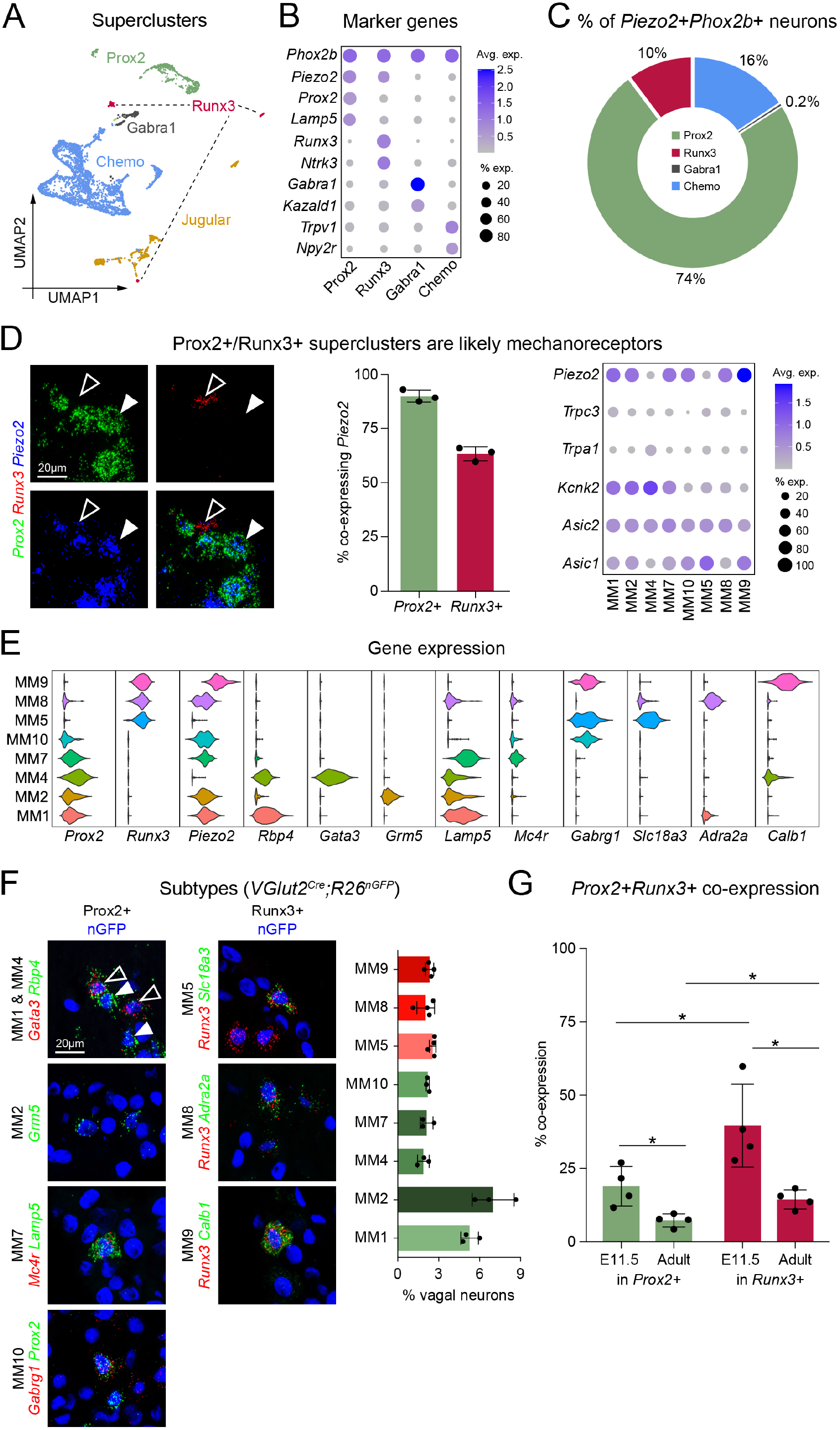
Meta-analysis of vagal neurons identified the Prox2+ and Runx3+ superclusters of putative mechanoreceptors. **(A)** Uniform manifold approximation and projection (UMAP) plot based on a meta-analysis of single cell RNA-sequencing data from this work and two published reports (Bai et al., 2019; Kupari et al., 2019), showing five vagal neuron superclusters (Prox2, Runx3, Chemo, Gabra1 and Jugular). **(B)** Dot plot showing the expression of selected genes that define the four Phox2b+ nodose superclusters (Prox2, Runx3, Gabra1 and Chemo). **(C)** Donut chart showing the supercluster identity of *Piezo2*+*Phox2b*+ nodose neurons. **(D) Left**, representative single molecule *in situ* fluorescent hybridization (smFISH) images for *Prox2* (green), *Runx3* (red) and *Piezo2* (blue) mRNA on vagal ganglia sections from wildtype mice at postnatal day (P) 8. **Middle**, quantifications showing that the majority of *Prox2*+ (90.0±2.8%, n = 3) and *Runx3*+ (63.8±3.2%, n = 3) neurons co-express *Piezo2*. **Right**, dot plot showing the expression of selected mechanosensory genes in the Prox2+/Runx3+ neuronal subtypes. Note that the Prox2+ MM4 and the Runx3+ MM5 subtypes are *Piezo2*-negative. **(E)** Violin plots showing the expression of selected genes that were used to identify the Prox2+/Runx3+ neuronal subtypes in panel F. **(F) Left**, smFISH images identifying Prox2+ neuronal subtypes MM1, MM2, MM4, MM7 and MM10. **Middle**, smFISH images identifying Runx3+ neuronal subtypes MM5, MM8, MM9. smFISH was performed on vagal ganglia sections from *VGlut2^Cre^;R26^nGFP^* mice at P8; vagal neuronal nuclei expressing nGFP were identified by immunofluorescence and are shown in blue. **Right**, quantification of Prox2+ and Runx3+ neuronal subtypes, n = 3 - 4. **(G)** Quantification of *Prox2+Runx3+* co-expression, n=4. Green bars show the percentage of *Prox2*+ neurons co-expressing *Runx3* at E11.5 (19.0±6.7%) and in the adult (7.3±2.2%), and red bars show the percentage of all Runx3+ neurons co-expressing Prox2 at E11.5 (39.6±14.2%) and in the adult (14.4±3.3%). The proportion of *Prox2*+*Runx3*+ double positive cells was greater at E11.5 than in adults both in the Prox2+ population (p = 0.0162) and in the Runx3+ population (p = 0.0134), and the proportion of *Prox2*+*Runx3*+ double positive cells was greater in the Runx3+ population than in the Prox2+ population both at E11.5 (p = 0.0388) and in the adult (p = 0.011). Data are represented as mean ± SD, *p < 0.05, unpaired two-tailed t-test. See also Figure S1.

A semi-supervised method was applied to group the 27 vagal neuronal subtypes into broader groups that were termed ‘superclusters’. The superclusters distinguish the two parts of the vagal ganglion, the nodose and jugular ganglion (also known as superior and inferior vagal ganglia), which derive from the epibranchial placode and cranial neural crest, respectively (Ayer-Le Lievre and Le Douarin, 1982; D’Amico-Martel and Noden, 1983). Whereas nodose neurons control the function of inner organs, jugular neurons allow somatic sensation and resemble dorsal root ganglia neurons. Nodose and jugular superclusters express *Phox2b* and *Prrxl1*, respectively, that are known to mark placodally-and neural crest-derived neuronal types (D’Autreaux et al., 2011; Pattyn et al., 1999) (Figure 1A and S1E,G). Three superclusters of nodose neurons were grouped together in the UMAP: ‘Prox2’ expressing the homeodomain transcription factor *Prox2*, ‘Gabra1’ expressing the gene encoding the alpha-1 (α1) subunit of the GABAA receptor and ‘Chemo’, a putative chemoreceptor supercluster expressing *TrpV1* and *Npy2r* (Figure 1A,B). The three remaining neuronal subtypes scattered in the UMAP, and all expressed the homeodomain transcription factor *Runx3;* they were merged into a fourth supercluster named ‘Runx3’. The Prox2 and Runx3 superclusters contained ∼84% of all nodose *Phox2b+Piezo2*+ neurons (Figure 1C, S1F,G). As Piezo2 transduces mechanical information in many neuron types (Coste et al., 2010; Murthy et al., 2017), we hypothesized that the Prox2 and Runx3 superclusters were candidate vagal mechanoreceptors innervating the internal organs. The remaining nodose *Phox2b+Piezo2*+ neurons were assigned to the ‘Chemo’ supercluster and might correspond to polymodal afferents (Figure 1C, S1F). Single molecule fluorescent *in situ* hybridization (smFISH) was used to examine the colocalization of *Piezo2*, *Prox2* and *Runx3* mRNA in vagal neurons (Figure 1D). In agreement with the sequencing data, we found that 90±3% of *Prox2*+ neurons and 63±3% of *Runx3*+ neurons co-expressed *Piezo2.* Prox2+ and Runx3+ subtypes express additional genes, such as *Kcnk2*, *Asic1/2*, *Trpc3* and *Trpa1* (Figure 1D) that have been implicated in mechanoreception (Ranade et al., 2015).

The Prox2+ and Runx3+ superclusters encompass five and three transcriptomically unique neuronal subtypes, respectively; we refer to these as *M*eta-analysis putative *M*echanoreceptors MM1, MM2, MM4, MM7, MM10 (Prox2+ subtypes) and MM5, MM8, MM9 (Runx3+ subtypes). Four and two of the Prox2+ and Runx3+ subtypes, respectively, express *Piezo2* (Figure 1D,E). Each individual subtype can be defined by the expression of a specific combination of marker genes, which was verified using smFISH on vagal ganglia sections from *VGlut2^Cre^;R26^nGFP^* mice at P8. Each subtype represents ∼2-8% of all vagal neurons (Figure 1F). In the adult mouse, the vast majority of Prox2 and Runx3 neurons exclusively expressed one or the other of these two transcription factors. In contrast, *Prox2* and *Runx3* are extensively co-expression during development. In particular, 40±14% of *Runx3*+ neurons co-expressed *Prox2* at E11.5, indicating that the Prox2+ and Runx3+ superclusters are developmentally related (Figure 1G).

### Prox2+/Runx3+ neurons form IGLEs in the upper gastrointestinal tract

We generated a *Prox2^FlpO^* knock-in transgenic mouse strain to further investigate putative vagal mechanoreceptors (Figure S2A). *In situ* hybridizations from the Allen Brain Atlas showed that *Prox2* expression is restricted to the cranial ganglia during development (Figure S2B), but in the adult *Prox2* is also expressed in other cell types (Nishijima and Ohtoshi, 2006). We therefore used an intersectional genetic strategy combining *Prox2^FlpO^* and *Phox2b^Cre^* with *Ai65*, a Tomato expressing reporter allele, to specifically label Prox2+ neurons in vagal ganglia (*Prox2^FlpO^;Phox2b^Cre^;Ai65* animals). Indeed, analysis of Tomato expression in adult mice showed that 95±4% of vagal *Prox2*+ neurons were recombined and expressed Tomato (Figure S2C). However, 71±2% of vagal *Runx3*+ neurons were also recombined, reflecting the shared developmental history of Prox2+/Runx3+ neurons (Figure S2C-G). Thus, recombination in *Prox2^FlpO^;Phox2b^Cre^;Ai65* animals occurs in both, Prox2+ and Runx3+ neurons, and we therefore refer to these mice hereafter as *Prox2/Runx3^Tom^*. To complement the histological analyses relying on the *Prox2/Runx3^Tom^* strain, the *Runx3^Cre^* allele was used. Runx3 is expressed in proprioceptive neurons of dorsal root ganglia and various non-neuronal cells types (Levanon et al., 2002; Shin et al., 2021). To restrict recombination to the vagal ganglia, we initially analyzed *Runx3^Cre^;Phox2b^FlpO^;Ai65* animals. Although vagal Runx3+ neurons were recombined in these mice, we occasionally observed recombination in enteric and sympathetic ganglia (data not shown). Therefore *Runx3^Cre^;Prox2^FlpO^;Ai65* animals were subsequently used to restrict recombination to Runx3+ vagal neurons (hereafter named *Runx3^Tom^* mice). Analysis of Tomato expression in *Runx3^Tom^* mice showed that 70±5% of vagal *Runx3*+ neurons were recombined (Figure S2D,H), but only a minor proportion of *Prox2+* neurons (Figure S2C). Thus, recombination occurred in all vagal Prox2+ and Runx3+ neuronal subtypes in *Prox2/Runx3^Tom^* mice, and was restricted to vagal Runx3+ neuronal subtypes in *Runx3^Tom^* mice.

We investigated the innervation sites of Prox2+/Runx3+ vagal neurons in the upper gastrointestinal tract using immuno-labeling of Tomato in *Prox2/Runx3^Tom^* and *Runx3^Tom^* mice. The esophagus and attached stomach were isolated, cleared and imaged using lightsheet microscopy (Susaki et al., 2015; Voigt et al., 2019). This demonstrated that the entire rostro-caudal axis of the esophagus was densely innervated by Prox2+/Runx3+ neurons, but only sparsely by Runx3+ vagal neurons (Figure 2A). Quantifications demonstrated that the vast majority of esophageal ganglia were contacted by Tomato+ IGLEs in *Prox2/Runx3^Tom^* mice, and rarely by Tomato+ IGLEs in *Runx3^Tom^* mice (Figure 2B). The axons tightly wrapped Phox2b+ enteric neurons, including both ChAT+ excitatory and Nos+ inhibitory enteric motor neurons (Figure 2C,D). Thus, our histological analyses indicated that the esophagus was mainly innervated by Prox2+ and, to a lesser extent, by Runx3+ neurons. Both, Prox2+ and Runx3+ neurons formed esophageal IGLEs.

**Figure 2.**
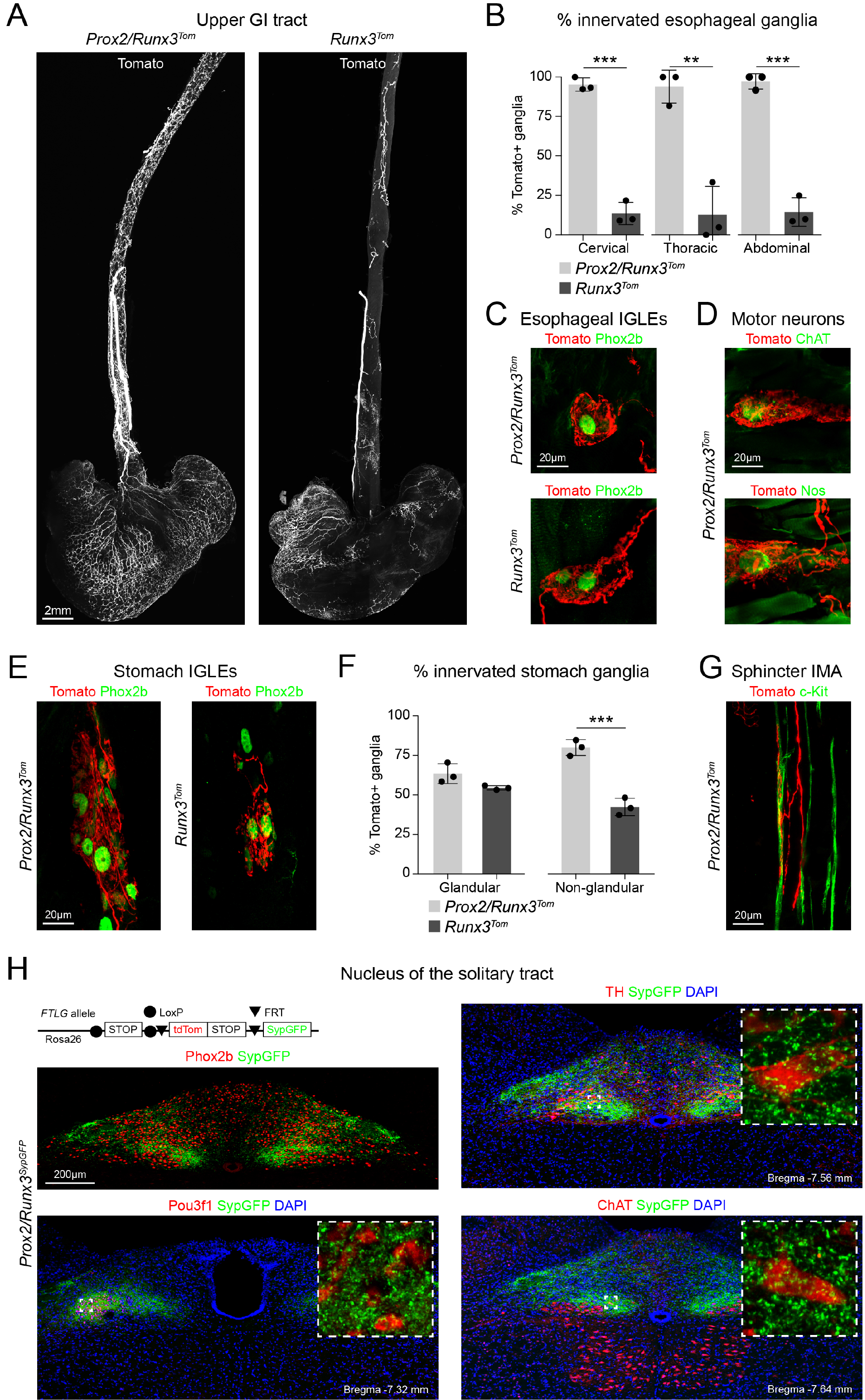
Prox2+/Runx3+ neurons form IGLEs and IMAs in the upper gastrointestinal tract. **(A)** Lightsheet imaging (acquired with a mesoSPIM) of the esophagus and stomach from *Prox2/Runx3^Tom^* mice (*Prox2^FlpO^;Phox2b^Cre^;Ai65)* **(left)** and *Runx3^Tom^* animals (*Runx3^Cre^;Prox2^FlpO^;Ai65*) **(right)** showing Tomato+ fibers (white) identified by immunohistology. **(B)** Quantification of the proportion of esophageal enteric ganglia, identified by Phox2b immunohistology, innervated by Tomato+ fibers in *Prox2/Runx3^Tom^* and *Runx3^Tom^* animals in the cervical (95.2±4.2% v. 13.6±7%, p = 0.0003), thoracic (93.9±10.5% v. 12.7±18.0%, p = 0.0053) and abdominal (97.2±4.8% v. 14.6±9.1%, p = 0.0007) esophagus, n = 3. **(C)** Immunohistological analysis of esophageal IGLEs in *Prox2/Runx3^Tom^* **(top)** and *Runx3^Tom^* animals **(bottom)** using antibodies against Tomato (red) and Phox2b (green). **(D)** Immunohistological analysis of innervated excitatory **(top)** and inhibitory **(bottom)** esophageal enteric motor neurons in *Prox2/Runx3^Tom^* animals using antibodies against Tomato (red), ChAT (green, top) and nNos (green, bottom). **(E)** Immunohistological analysis of stomach IGLEs in *Prox2/Runx3^Tom^* **(left)** and *Runx3^Tom^* mice **(right)** using antibodies against Tomato (red) and Phox2b (green). **(F)** Quantification of the proportion of stomach enteric ganglia, identified by Phox2b immunohistology, innervated by Tomato+ fibers in *Prox2/Runx3^Tom^* and *Runx3^Tom^* animals in the glandular (63.5±6.3% v. 54.6±1.3%, p = 0.127) and non-glandular (80.1±5.1% v. 42.5±5.5%, p = 0.001) stomach, n = 3. **(G)** Immunohistological analysis of IMAs in *Prox2/Runx3^Tom^* animals using antibodies against Tomato (red) and c-Kit (green). **(H)** Analysis of the central synaptic connectivity of Prox2+/ Runx3+ neurons in the nucleus of the solitary tract. Scheme **(top left)** of the intersectional *FTLG* reporter allele used to label Prox2+ and Runx3+ synapses with SypGFP in *Prox2/Runx3^SypGFP^* (*Prox2^FlpO^;Phox2b^Cre^;R26^FTLG^*) mice (Dempsey et al., 2021). Immunohistological analysis of animals showing dense GFP+ boutons (green) in the NTS **(top left)**, on Pou3f1+ (red, **bottom left**) and TH+ (red, **top right**) NTS neuronal subtypes, as well as on ChAT+ neurons (red, **bottom right**) of the dorsal motor nucleus of the vagus. Data in **(B, F)** are represented as mean ± SD, **p < 0.01, ***p < 0.001, unpaired two-tailed t-test. See also Figure S2.

In the stomach, we detected Tomato+ IGLEs in the glandular and non-glandular regions in *Prox2/Runx3^Tom^* and in *Runx3^Tom^* animals (Figure 2A,E). The lightsheet analysis of the *Runx3^Tom^* animal showed a higher density of innervation in the glandular than the non-glandular stomach (Figure 2A,F). Thus, both Prox2+ and Runx3+ neurons form gastric IGLEs, but the Prox2+ neurons preferentially innervate the non-glandular stomach and the Runx3+ neurons the glandular stomach. Tomato+ axons also made contacts with c-Kit+ interstitial cells of Cajal in the esophageal sphincter of *Prox2/Runx3^Tom^*, but not *Runx3^Tom^* mice; these endings correspond to IMAs (Figure 2G). Occasional Tomato+ IMAs were detected in the stomachs of both *Prox2/Runx3^Tom^* and *Runx3^Tom^* mice, but these were rare.

### Central projections of Prox2+/Runx3+ neurons

To define the synaptic connectivity of Prox2+/Runx3+ vagal neurons to second order sensory neurons in the nucleus of the solitary tract, we labeled their synapses with SynaptophysinGFP using *Prox2/Runx3^SypGFP^*(*Prox2^FlpO^;Phox2b^Cre^;R26^FTLG^* mice (Dempsey et al., 2021); see Figure 2H for a schema of the FTLG allele). This approach revealed many GFP+ synaptic boutons on Phox2b+ neurons in the nucleus of the solitary tract (NTS) (Figure 2H). Pou3f1+ and TH+ neurons in the NTS represent small populations of interneurons found in the central and ventral nuclei of the NTS, respectively, that were innervated by Prox2+/Runx3+ vagal sensory neurons. Interestingly, we also found GFP+ boutons on ChAT+ neurons of the dorsal motor nucleus of the vagus. Thus, Prox2+/Runx3+ vagal neurons project to Pou3f1+ and TH+ neurons in the NTS, and to directly to ChAT+ motorneurons in the dorsal motor nucleus of the vagus.

### Prox2+/Runx3+ neurons innervate the upper gastrointestinal tract

Next, we aimed to identify putative mechanoreceptive neurons that innervate the upper gastrointestinal tract. We retrogradely labeled vagal sensory neurons projecting to various sites in the esophagus and stomach (cervical, thoracic and abdominal esophagus, glandular and non-glandular stomach; shown schematically in Figure 3A) by injections of Cholera Toxin b (CTb) and Fast Blue (Bentivoglio et al., 1980; Stoeckel et al., 1977). smFISH on vagal ganglia sections was used to define *Prox2, Runx3* and *Piezo2* expression in the retrogradely traced neurons (Figure 3B). A higher proportion of vagal neurons that innervate the esophagus were *Piezo2+* as compared to those that innervate the stomach (Figure S3A). Overall, most traced *Piezo2*+ neurons from the upper gastrointestinal tract corresponded to Prox2+ or Runx3+ neuronal subtypes, i.e. ∼75% from the cervical and thoracic esophagus and ∼100% from the abdominal esophagus and stomach (Figure 3B). The upper esophagus is innervated by both jugular and nodose neurons (Kwong et al., 2008), and we suggest that jugular neurons represent the remaining back-traced *Piezo2*+ cells.

**Figure 3.**
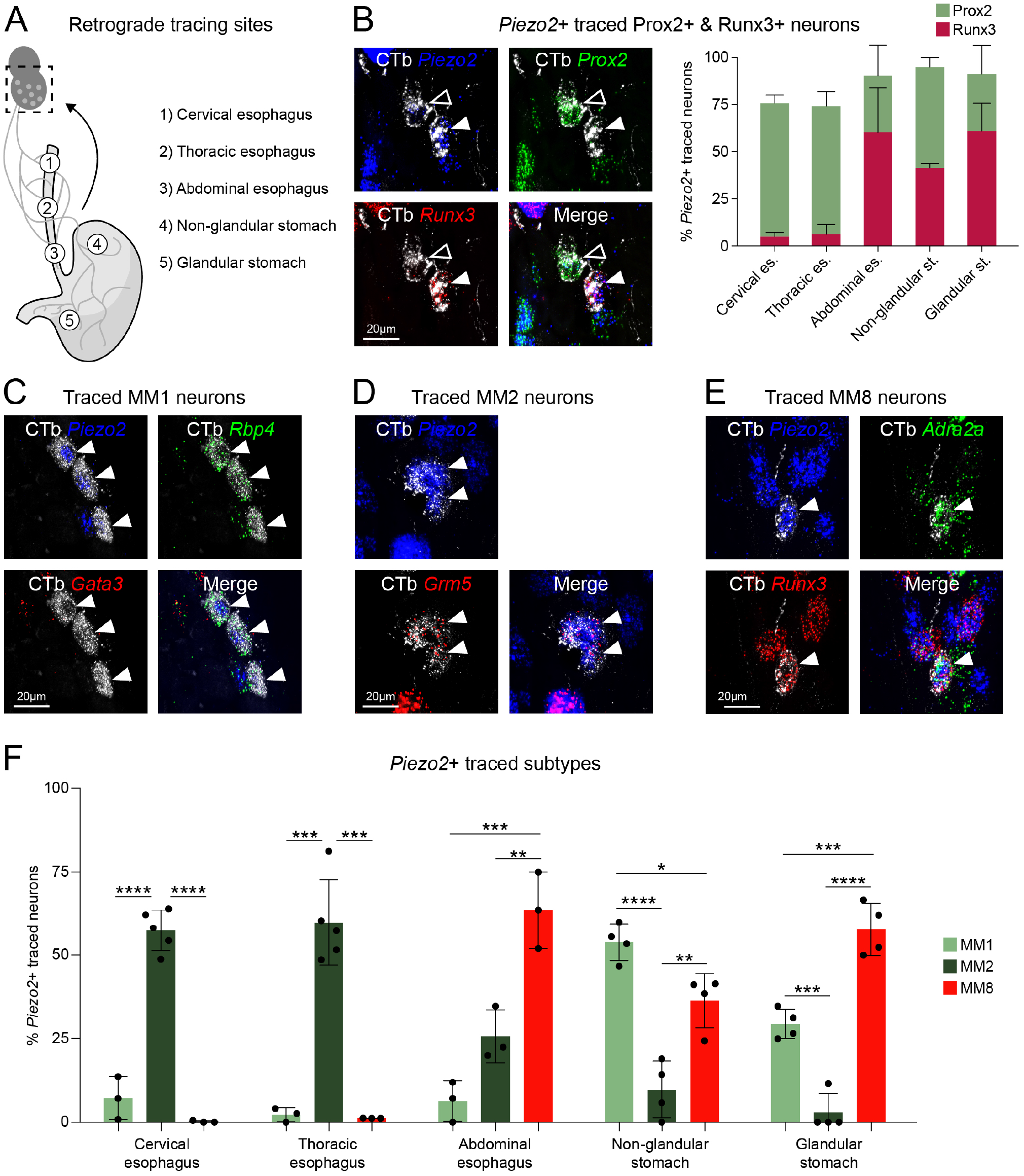
MM1, MM2 and MM8 neurons are the major subtypes of Prox2+/Runx3+ vagal neurons innervating the upper digestive tract. **(A)** Retrograde tracing experiments; the scheme shows the five injection sites analyzed: 1 cervical esophagus, 2 thoracic esophagus, 3 abdominal esophagus, 4 non-glandular stomach, 5 glandular stomach. **(B) Left**, smFISH analysis of neurons after injection of CTb into the abdominal esophagus; probes used were *Prox2* (green), *Runx3* (red) and *Piezo2* (blue), which was combined with immunohistological analysis of CTb (gray). The open arrowhead marks a *Prox2*+*Piezo2*+CTb+ cell, and the filled arrowhead marks a *Runx3*+*Piezo2*+CTb+ cell. The proportions of the *Prox2*+*Piezo2*+ and *Runx3*+*Piezo2*+ neurons that were retrogradely labeled from the different injection sites are quantified on the **right**. Note that for technical reasons we used Fast Blue to trace from the cervical and thoracic esophagus in postnatal P7 mice, and CTb to trace from the abdominal esophagus and stomach in adult mice (see Methods for more details, n = 3 - 5). Runx3+ neurons corresponded to 5.1±2.1%, 6.3±5.1%, 60.3±23.5%, 41.5±2.5%, and 61.1±14.6% of all *Piezo2*+ traced neurons traced from the cervical, thoracic, and abdominal esophagus, and from the non-glandular and glandular stomach, respectively. Prox2+ neurons corresponded to 70.6±4.4%, 67.8±7.6%, 29.8±16.2%, 53.4±5.2%, and 30.1±15.2% of all *Piezo2+* neurons traced from the cervical, thoracic, and abdominal esophagus, and from the non-glandular and glandular stomach, respectively. **(C-E)** smFISH analysis of traced MM1, MM2 and MM8 neurons. CTb (gray) was detected by immunohistology. For the identification of MM1 neurons (C), probes against *Rbp4* (green), *Gata3* (red, negative marker) and *Piezo2* (blue) mRNA. To identify MM2 neurons (D), probes against *Grm5* (green) and *Piezo2* (blue) mRNA were used. To identify MM8 neurons (E), probes against *Adra2a* (green), *Runx3* (red) and *Piezo2* (blue) mRNA were used. **(F)** Quantification of traced MM1, MM2 and MM8 neurons from the five injection sites (n = 3 - 5). MM2 neurons were the dominant subtype traced from the cervical and thoracic esophagus (57.5±6.1% and 59.9±12.8%), whereas MM1 (7.2±6.4% and 2.3±2.1%) and MM8 (0.2±0.3% and 1.3±0.03%) were rarely traced’. The abdominal esophagus was largely innervated by MM8 neurons (63.6±11.5%), with MM2 neurons contributing a substantial proportion (25.8±7.9%) and MM1 neurons contributing a small proportion (6.4±6.0%). The non-glandular stomach was mostly innervated by MM1 neurons (53.93±5.6%), with a large contribution of MM8 neurons (36.4±8.1%) and a small contribution of MM2 neurons (9.8±8.5%). The glandular stomach was largely innervated by MM8 neurons (57.8±7.9%), with a large contribution of MM1 (29.4±4.4%) and a small contribution of MM2 neurons (2.9±5.8%). Data are represented as mean ± SD, *p < 0.05, **p < 0.01, ***p < 0.001, ****p < 0.0001, Ordinary one-way ANOVA with Tukey’s multiple comparisons test. See also Figure S3.

Further analysis demonstrated that the Prox2+ MM1 and MM2, as well as the Runx3+ MM8 neuronal subtypes correspond to the *Piezo2*+ neuronal subtypes innervating the esophagus and stomach (Figure 3C,D,E). These three subtypes displayed preferences in their regional innervation patterns. The cervical and thoracic esophagus were mainly innervated by the Prox2+ MM2 neuronal subtype. MM2 neurons were less frequently traced from the abdominal esophagus and rarely from the stomach (Figure 3D,F). Instead, the related Prox2+ neuronal subtype MM1 was more frequently traced from the stomach (Figure 3C,F). The Runx3+ neuronal subtype MM8 was rarely traced from the cervical and thoracic esophagus, but more frequently from the abdominal esophagus and stomach (Figure 3E,F). We detected further regional differences in the stomach: MM8 and MM1 were more frequently traced from the glandular and non-glandular stomach, respectively (Figure 3F). Previous work showed that *Glp1r*+ vagal neurons form gastric IGLEs (Williams et al., 2016). smFISH revealed extensive *Glp1r* expression in MM1 (89.6±5.3%) and MM8 (94.3±2.6%) neurons, but the esophageal projecting MM2 neurons rarely expressed *Glp1r* (2.4±3.0%) (Figure S3B,C). In summary, retrograde tracing and histological analyses show that the MM1, MM2 and MM8 subtypes innervate the esophagus and stomach where they end as IGLEs. These subtypes display regional innervation preferences and, in particular, the esophagus is mainly innervated by the Prox2+ MM2 subtype.

### Prox2+/Runx3+ vagal neurons function as mechanoreceptors

We next investigated the electrophysiological properties of Prox2+/Runx3+ neurons using an *in vitro* open book vagus-esophagus preparation (Figure 4A) to record mechanically evoked responses from single nerve fibers that innervate the thoracic and abdominal esophagus (Page and Blackshaw, 1998; Page et al., 2002). For this, *Prox2/Runx3^ChR^*(*Prox2^FlpO^;Phox2b^Cre^;Ai80*) mice that express channelrhodopsin specifically in Prox2+/Runx3+ vagal neurons were used (Daigle et al., 2018). Neurons were classified as Prox2/Runx3-positive or -negative based on whether or not they fired action potentials in response to blue light (Figure 4B and Figure S4A,B). Firing patterns evoked by light and mechanical stimuli had similar magnitude and kinetics (Figure 4B). Interestingly, all mechanically sensitive neurons tested in the thoracic and abdominal esophagus responded to blue light (30/30 neurons), indicating that Prox2/Runx3-positive neurons are the sole vagal mechanoreceptors at this site (Figure 4C,D). Prox2/Runx3-positive neurons were responsive to forces ranging from 60 to 250 mN delivered by a piezo actuator and increased their firing frequency in response to increasing force (Figure 4B,E). Analysis of single unit responses to mechanical stimuli revealed two distinct response patterns. The majority of the Prox2/Runx3-positive mechanoreceptors (80%) showed sustained firing of up to 20Hz during stimulation, and firing frequency slowly returned to baseline after the end of the stimulation (we call these type I esophageal receptors, Figure 4F). The remaining 20% showed peak firing of around 7Hz that rapidly returned to baseline before the end of the stimulus (we call these type II esophageal receptors, Figure 4F). In summary, Prox2/Runx3-positive neurons form esophageal IGLEs and function as low-threshold mechanoreceptors. These neurons segregate into an abundant type I and a less abundant type II receptor that differ in their adaptation rates, and we suggest that these correspond to MM2 and MM8, respectively.

**Figure 4.**
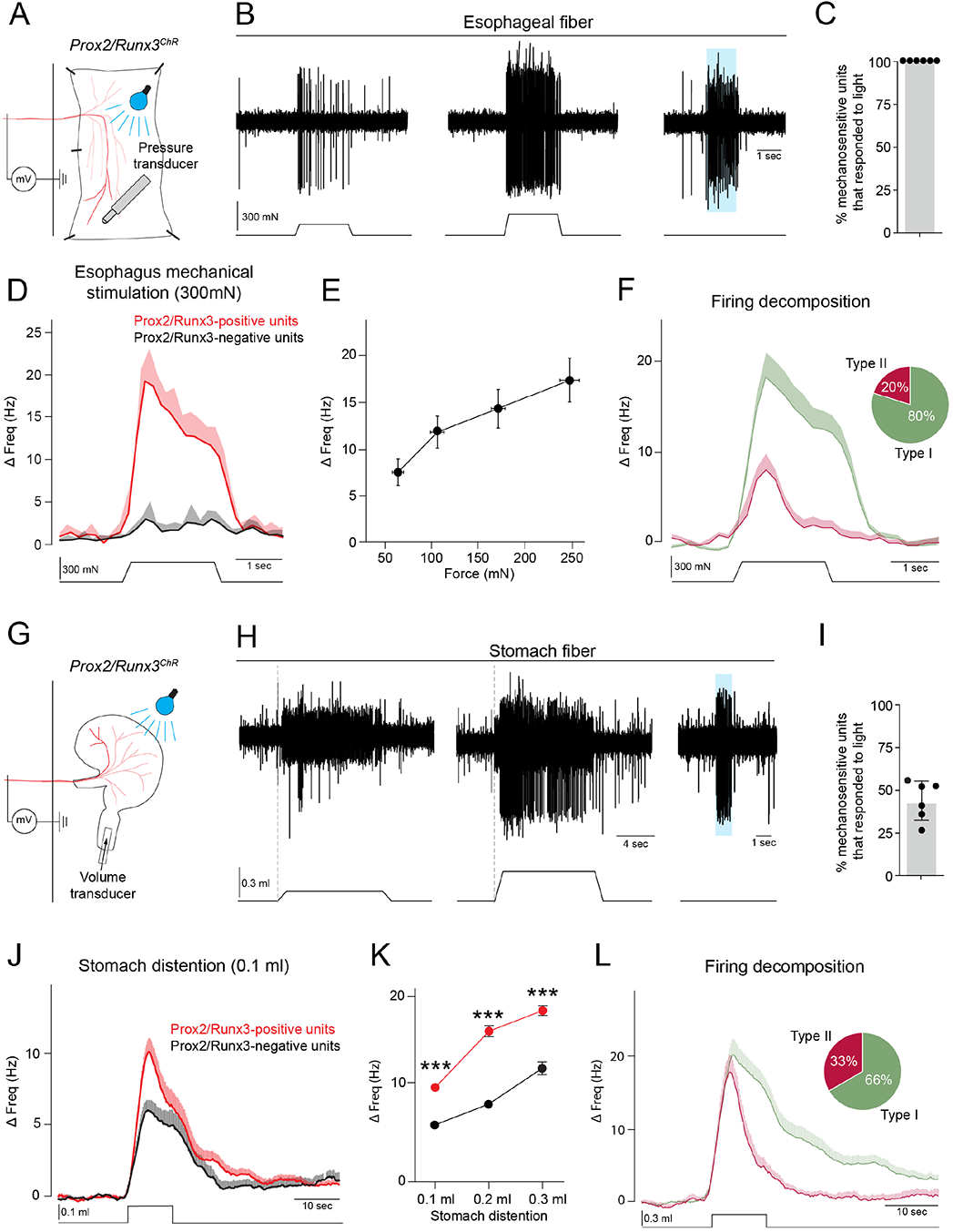
Prox2+/Runx3+ neurons are esophageal and gastric mechanoreceptors. **(A)** Scheme of the open book esophagus-vagus nerve *in vitro* preparation used for electrophysiological analysis. Teased fibers from the vagus nerve of *Prox2/Runx3^ChR^* (*Prox2^FlpO^;Phox2b^Cre^;Ai80*) mice were placed on a recording electrode; the identity of Prox2/Runx3-positive units expressing channelrhodopsin was identified by light stimulation. Esophageal mechanosensitive receptive fields were found by probing the tissue with a blunt glass rod and then a piezo actuator. **(B)** Representative traces of a vagus nerve fiber responding to mechanical stimuli of increasing strength **(top)**; the intensity of the mechanical stimulus is indicated below (**Left**, 100 mN; **middle**, 250 mN; **right**, 0 mN). Note that this fiber also responded to blue light applied to the receptive field, demonstrating that it contained Prox2/Runx3-positive units. **(C)** Quantification of the proportion of Prox2/Runx3-positive units (30/30) among all mechanosensitive units in the esophagus; note that all mechanosensitive fibers were Prox2/Runx3-positive units, n = 6 mice. **(D)** Quantification of population firing patterns of Prox2/Runx3-positive (red) and – negative (black) units during strong mechanical stimulation (300mN). In the esophagus, only Prox2/Runx3-positive units (red) responded to mechanical stimulation. **(E)** Stimulus response properties of Prox2/Runx3-positive units to increasing mechanical force (63.3±6.0 – 247.4±10.5 mN, n = 31 units). **(F)** Decomposition of Prox2/Runx3-positive unit firing patterns revealed two types of mechanosensory neurons that differed in their adaptation properties (n = 30 units). The firing of 80% of the recorded Prox2/Runx3-positive units decayed after the end of the stimulus (type I, green, n = 24), whereas in the remaining 20% firing decayed during the stimulus (type II, red, n = 6). **(G)** Scheme of the stomach-vagus nerve *in vitro* preparation. In order to mimic physiological distension, the stomach was kept intact and a catheter was inserted into the pyloric sphincter. Stomach distension was induced by injecting 0.1, 0.2 or 0.3 ml of a saline solution through the catheter into the stomach. **(H)** Traces recorded from a vagus nerve fiber responding to increasing distention stimuli, i.e. 0.1 ml and 0.3 ml stimuli, and to light stimulation. **(I)** Quantification of the proportion of Prox2/Runx3-positive units among all mechanosensitive units in the stomach (55/125, n = 6 mice). **(J)** Population firing responses of Prox2/Runx3-positive (n = 54) and -negative (n = 36) units during stomach distention (0.1 ml). **(K)** Stimulus response properties of Prox2/Runx3-positive (n = 45-54) and -negative (n = 35-43) units to increasing stomach distension (0.1 – 0.3 ml). **(L)** Decomposition of Prox2/Runx3-positive unit firing patterns revealed two types (n = 46 units). All units responded rapidly to stomach distention, but differed in how their firing adaptation. Type I units (66%, green, n = 31) were slowly adapting units that returned to basal activity after stimulus offset. Firing of type II units (33%, red, n = 15) decayed during the duration of the stimulus. Data are represented as mean ± SD **(C,I)** or mean ± SEM **(D,E,F,J,K,L)**, ***p < 0.001, unpaired two-tailed t-test. See also Figure S4.

In addition, we used an *in vitro* vagus-stomach preparation (Figure 4G), and administered physiological volumes of saline (0.1-0.3ml) as a mechanical stimulus to distend the stomach (Kim et al., 2020a; Williams et al., 2016). Around half of all distention-sensitive neurons were light-sensitive and thus corresponded to Prox2/Runx3-positive neurons (55/125 neurons; Figure 4H-K). Examples of mechanosensitive single units, including both light-sensitive and -insensitive receptors, are shown in Figure S4C. Across all volumes tested, light-sensitive neurons displayed higher firing rates compared to light-insensitive ones (Figure 4J,K). Closer examination of Prox2/Runx3-positive mechanosensitive firing responses uncovered two distinct types (Figure 4L). In both firing frequency increased at the onset of stomach distension, but they differed in their adaptation properties. Type I gastric receptors continued firing while the stomach was distended and only returned to basal activity after the stomach had emptied, while type II gastric receptors reduced their firing frequency and returned to basal activity even while the stomach was still distended (Figure 4L). Together, our *in vitro* studies demonstrate that Prox2/Runx3-positive gastric vagal neurons are mechanosensitive. Type I gastric receptors appear to detect static stretch or volume, and type II receptors the dynamic changes in stretch or tension. We propose that type I and II gastric receptors correspond to MM1 and MM8, respectively (see Discussion).

### Ablation of Prox2+/Runx3+ neurons results in esophageal dysmotility

We used a similar intersectional genetic strategy as the one described above to express the diphtheria toxin receptor (DTR) in Prox2+/Runx3+ neurons (*Prox2^FlpO^;Phox2b^Cre^;Tau^ds-DTR^*, hereafter called *Prox2/Runx3^ds-DTR^* mice) (Britz et al., 2015). DTR expressing neurons were ablated by injecting diphtheria toxin (DT), and the ablation efficacy and specificity was determined using smFISH two weeks after DT injection. As the DTR receptor is only expressed after the removal of both lox-flanked and frt-flanked stop cassettes, DT-treated *Prox2^FlpO^;Tau^ds-DTR^* animals were used as controls (called *Control^ds-DTR^*). This showed that 97±3% and 90±5% of *Prox2*+ and *Runx3*+ neurons, respectively, were ablated in the vagal ganglia of *Prox2/Runx3^ds-DTR^* animals (Figure 5A). The scRNAseq data showed that Prox2+ and Runx3+ neurons represent around 30% of all Phox2b+ vagal neurons. In accordance, 32±6% of Phox2b+ neurons were ablated after DT injection (Figure 5B). Following administration of DT, the animal’s weight was monitored daily, which revealed that *Prox2/Runx3^ds-DTR^* but not *Control^ds-DTR^* mice rapidly lost weight in the first five days post ablation (Figure 5C and S5A). *Prox2/Runx3^ds-DTR^* animals received a high caloric diet and daily saline injections after ablation, which helped to stabilize their weight at about −12% of the starting weight (Figure 5C). We performed videofluoroscopic swallowing studies (VFSS) 2 days before and 19 days after ablation (Figure 5D,E and Supplemental Videos 1, 2, see Figure S5B for an experimental timeline) (Lever et al., 2015). The esophageal transit time of a liquid bolus increased from an average of 1.5±0.4 seconds to 39.3±15.4 seconds after ablation (Figure 5F). Additionally, the liquid bolus was frequently retained in the esophagus, flowed in an orad direction to re-enter the pharynx which in some cases resulted in projectile regurgitation from the mouth (Supplemental Video 3). These aberrant esophageal bolus flow behaviors were never observed before ablation. This was accompanied by megaesophagus, with the average diameter of their abdominal esophagus increasing from an average of 1.7±0.2 mm before ablation to 2.8±0.2 mm after ablation (Figure 5F). Moreover, numerous examples of aerophagia were observed after ablation (Supplemental Video 4). In addition, pharyngeal transit time was increased, whereas lick rate and lick time were slightly decreased and increased, respectively (Figure 5F). All other parameters related to the biomechanics of swallowing (swallow rate, swallow interval, lick to swallow ratio, jaw opening/closing velocity) were unaffected (Figure S5C). In summary, we observed severe deficits in ingestion after the ablation of Prox2+/Runx3+ neurons that were assigned to esophageal dysmotility.

**Figure 5.**
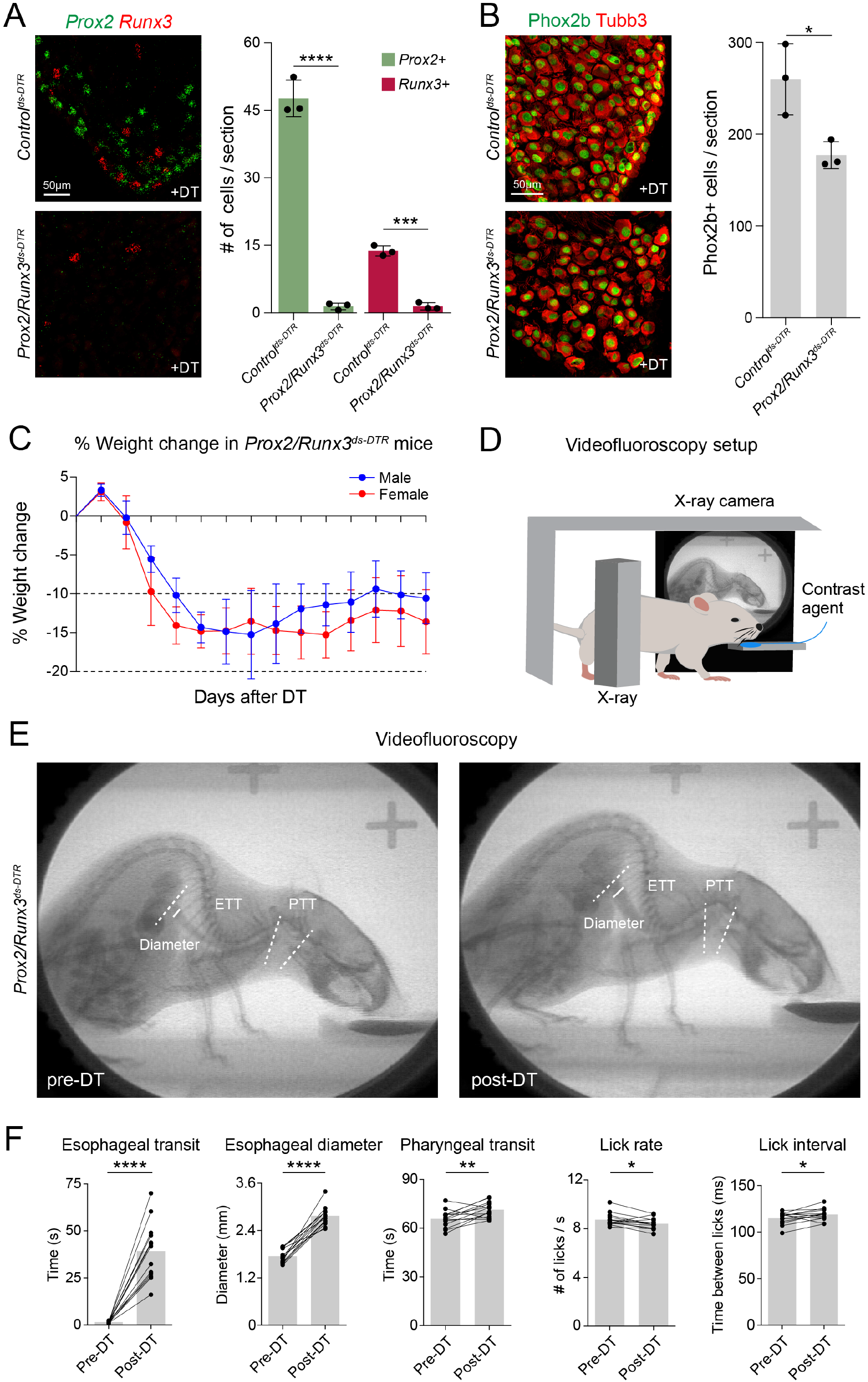
Ablation of Prox2+/Runx3+ neurons impairs esophageal motility in freely behaving animals. **(A)** Representative smFISH images **(left)** against *Prox2* (green) and *Runx3* (red) mRNA in adult *Control^ds-DTR^* (**top**, *Prox2^FlpO^;Tau^ds-DTR^*) and *Prox2/Runx3^ds-DTR^* (**bottom**, *Prox2^FlpO^;Phox2b^Cre^;Tau^ds-DTR^*) mice 14 days after DT administration. The number of *Prox2*+ and *Runx3*+ neurons before and after ablation are quantified on the right, n = 3. The number of *Prox2*+ neurons per section decreased from 47.6±4.1 in *Control^ds-DTR^* to 1.5±0.7 in *Prox2/Runx3^ds-DTR^* mice (p < 0.0001), while the number of *Runx3*+ neurons per section decreased from 13.8±1.1 in *Control^ds-DTR^* to 1.5±0.8 in *Prox2/Runx3^ds-DTR^* mice (p = 0.0001). **(B)** Representative immunofluorescence images **(left)** against Phox2b (green) and Tubb3 (red) in adult *Control^ds-DTR^* (**top**) and *Prox2/Runx3^ds-DTR^* (**bottom**) mice 14 days after DT administration. The number of Phox2b+ neurons before and after ablation are quantified on the right, n = 3. The number of Phox2b+ neurons per section was 260.0±38.8 in *Control^ds-DTR^* and 177.2±14.7 in *Prox2/Runx3^ds-DTR^* mice (p = 0.0259). **(C)** Graph showing the weight change in *Prox2/Runx3^ds-DTR^* mice for 14 days following DT administration, n = 14. **(D)** Scheme of the videofluoroscopy setup. **(E)** Single frame X-ray images from videofluoroscopy videos taken from one *Prox2/Runx3^ds-DTR^* mouse before (pre-DT, **left**) and after (post-DT, **right**) DT administration. Dotted white lines show the bolus traveling distance used to determine pharyngeal transit time (PTT) and esophageal transit time (ETT). Solid white line shows the site used to measure the esophageal diameter. **(F)** Esophageal transit time increased from an average of 1.5±0.5 s to 39.3±15.4 s after ablation, p < 0.0001. Esophageal diameter increased from an average of 1.7±0.2 mm to 2.8±0.2 mm after ablation, p < 0.0001. Pharyngeal transit time increased from an average of 65.8±5.3 ms to 71.3±4.8 ms after ablation, p = 0.0051. Lick rate decreased from an average of 8.7±0.5 licks/s to 8.4±0.5 licks/s after ablation, p = 0.0176. Lick interval increased from an average of 114.9±6.3 ms to 118.9±6.4 ms after ablation, p = 0.0335, n = 13 - 14. Data are represented as mean ± SD, *p < 0.05, ***p < 0.001, ****p < 0.0001, unpaired two-tailed t-test **(A,B)**, paired two-tailed t-test **(F)**. See also Figure S5.

## Discussion

Sensory neurons of the vagus nerve survey the mechanical state of the gastrointestinal tract and monitor esophageal and gastric distension. Here we used genetically guided anatomical tracing and optogenetic tools to show that three vagal sensory neuronal subtypes that express *Prox2* and *Runx3* innervate the esophagus and stomach with regionalized specificity. All three subtypes form IGLEs on enteric ganglia and function as low threshold mechanoreceptors, but they display different adaptation properties (See Figure 6 for a summary). We genetically ablated Prox2+/Runx3+ neurons, and demonstrate that this resulted in dysphagia due to severe esophageal dysmotility in freely behaving animals. Our results reveal the importance of vagal sensory feedback provided by Prox2+/Runx3+ neurons in swallowing and food intake.

**Figure 6.**
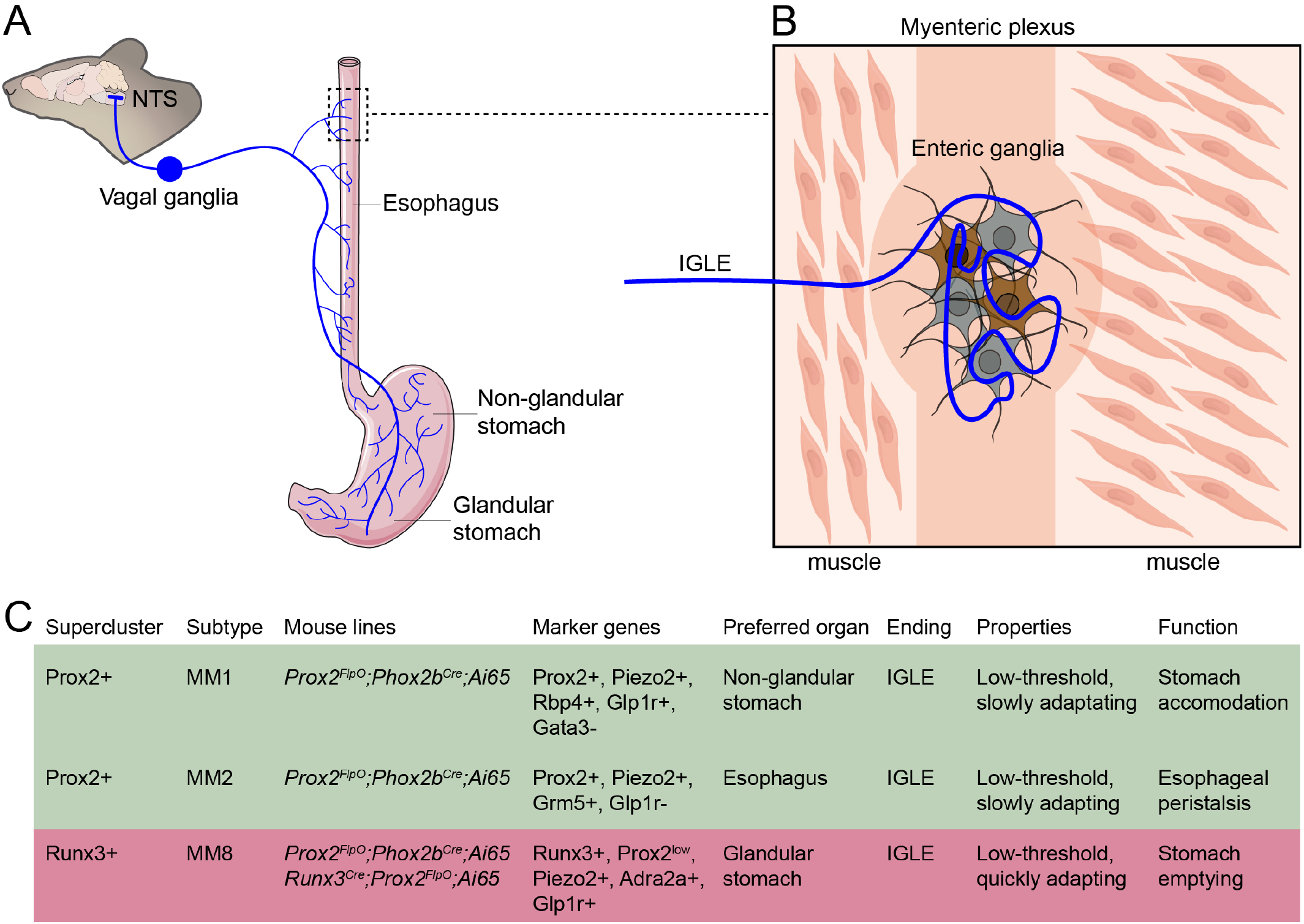
Summary. **(A)** Scheme of MM1, MM2 and MM8 vagal neurons and their targets in the upper gastrointestinal tract. Illustrations were adapted from bioicons.com (digestive-system-exploded, Servier; smooth-muscle-fiber, Servier; neuron, DBCLS) and scidraw.io (Mouse Brain Silhouette, Ann Kennedy) and licensed under CC-BY 3.0 and CC-BY 4.0. **(B)** Scheme of an MM2 IGLE contacting an esophageal enteric ganglion. Note that both excitatory (gray) and inhibitory (brown) esophageal motor neurons are contacted by the IGLE. **(C)** Table summarizing the characteristics of MM1, MM2 and MM8 vagal neurons. Indicated are the mouse lines used to label the neuronal subtypes, marker genes used to identify them, the preferred organ innervated by the neuronal subtypes, their mechanosensitive properties, and their putative functions in digestive physiology.

### Prox2+/Runx3+ neurons that innervate the esophagus and stomach

Neurons of the Prox2+/Runx3+ lineage represent the majority (∼85%) of all *Piezo2*+ neurons in the nodose ganglion. This lineage encompasses eight neuronal subtypes and three of these, the MM1, MM2 and MM8 subtypes, form IGLEs that innervate enteric ganglia in the esophagus and stomach.

Together, MM2 and MM8 neurons of the Prox2+/Runx3+ lineage innervate almost all (>95%) esophageal enteric ganglia, with MM2 being by far the more abundant subtype. Despite the morphological similarity of the nerve endings, we were able to distinguish between MM2 and MM8 neurons in our genetically guided anatomical tracing experiments. The ablation of the Prox2+/Runx3+ neurons demonstrated their importance in swallowing physiology. Specifically, their ablation resulted in a dramatic increase in the transit time of a bolus through the esophagus accompanied by megaesophagus. Esophageal peristalsis is part of a complex motor sequence involving the consecutive and stereotypic contractions of pharyngeal and esophageal muscle groups. This intricate sequence of events is controlled by a central pattern generator in the brainstem (Jean, 2001). Deafferentation of the thoracic esophagus in sheep provided early evidence of the importance of vagal sensory feedback in esophageal peristalsis, but the molecular nature of the vagal sensory neurons providing the essential signals has remained elusive (Falempin *et al*., 1986). Our results show that the MM2 and MM8 neuronal subtypes of the Prox2/Runx3 lineage provide the necessary peripheral feedback for esophageal peristalsis.

Previous work showed that *Glp1r* labels neurons that form gastric IGLEs (Williams *et al*., 2016), and we show that MM1 and MM8 both express *Glp1r*, indicating previously unappreciated heterogeneity in gastric IGLEs. In the esophagus, IGLEs are formed by MM2 and to a lesser extent by MM8 neuronal subtypes, but the MM2 subtype is Glp1r-negative. Others previously described an Oxtr+ subtype of vagal sensory neurons that forms IGLEs on intestinal ganglia (Bai *et al*., 2019). Our meta-analysis revealed significant *Oxtr* expression in 8 clusters, including MM2. Thus, despite their apparent morphological uniformity, IGLEs in the gastroenteric tract originate from molecularly heterogeneous neuron subtypes.

### Electrophysiological properties of Prox2+/Runx3+ neurons innervating the upper gastrointestinal tract

Previous analyses of gastrointestinal mechanoreceptors distinguished electrophysiological responses to stretch, tension and mechanical probing, uncovering both muscular and mucosal mechanoreceptors in various species (Kim *et al*., 2022; Page and Blackshaw, 1998; Page *et al*., 2002; Zagorodnyuk and Brookes, 2000; Zagorodnyuk *et al*., 2003). The use of optogenetic tools allowed us to analyze a small subset of morphologically and molecularly defined neurons and to determine their electrophysiological properties. Prox2+/Runx3+ neurons represent all vagal *Piezo2*+ neurons innervating the stomach. Nevertheless, only about half of all mechanoreceptors that respond to stomach distention correspond to Prox2+/Runx3+ neurons, demonstrating that Piezo2-negative gastric mechanoreceptors exist. However, Prox2+/ Runx3+ mechanoreceptors responded with higher firing frequencies to distention stimuli than the other types of gastric mechanoreceptors that do not express Piezo2.

In accordance with the molecular and anatomical data, our electrophysiological experiments distinguished different mechanoreceptive subtypes of the Prox2/Runx3 lineage. The esophageal type I and type II low threshold mechanoreceptors possess distinct adaptation properties, with type I receptors adapting more slowly than the type II receptors. Type I outnumbered type II mechanoreceptors four to one. MM2 are the abundant Prox2/Runx3 neuronal subtype innervating the esophagus. We therefore propose that MM2 neurons correspond to esophageal type I mechanoreceptors, and MM8 to type II receptors. It is possible that these neurons encode distinct aspects of esophageal distension. Esophageal distension mediates reflexes that result in contraction of the upper esophageal sphincter, and these differentially respond to slow and fast esophageal distention (Lang, 2009; Lang et al., 2001). Here, we analyzed the consequences of simultaneously ablating MM2 and MM8 subtypes, but it would be interesting to dissect the relative contributions of the two different subtypes to esophageal reflexes in future studies.

Prox2+/Runx3+ mechanoreceptors in the stomach also differed in their adaptation properties. Both, gastric and esophageal type I receptors displayed similar properties and were slowly adapting. We propose that the gastric type I receptors correspond to MM1, whereas we assign the esophageal type I receptor to the transcriptionally related MM2 subtype. Further, esophageal and gastric type II receptors display similar adaption properties, and we suggest that these correspond to the MM8 subtype. A recent study recorded calcium activity in vagal neurons during stomach/esophagus distension and found that Group C neurons (in particular *Piezo2+Glp1r+Rbp4+ and Piezo2+Grm5+Slit2+* neurons that appear to correspond to our MM1 and MM2 subtypes) display a slowly adapting response, similar to our type I mechanoreceptors (Zhao *et al*., 2022).

The slowly adapting MM1 and rapidly adapting MM8 gastric IGLE subtypes are regionally specialized, preferentially innervating the non-glandular and glandular stomach, respectively. These portions of the stomach have different functional roles during digestion (Janssen et al., 2011). During feeding, the ingested material collects in the non-glandular stomach that acts as a food reservoir and slowly expands in a process known as gastric accommodation. Fast peristaltic contractions then mechanically break down the ingested material in the glandular stomach (Janssen *et al*., 2011). Based on their regional specificity and functional responses, we hypothesize that the slowly adapting MM1 type I mechanoreceptors specialize to detect the volume changes that occur in the non-glandular stomach during gastric accommodation, while the fast adapting MM8 type II mechanoreceptors detect phasic stretch that accompany peristaltic contractions in the glandular stomach.

### Neuronal control of esophageal motility

Abnormal esophageal motility encompasses a heterogeneous class of disorders of frequently unclear etiology (Kahrilas et al., 2015). Hereditary, autoimmune, and infectious factors, as well as nervous system degeneration are all suggested to cause or contribute to this pathology (Boeckxstaens et al., 2014). In humans, videofluoroscopy swallow studies (VFSS) are used to diagnose and monitor swallowing dysfunction (Martin-Harris and Jones, 2008). Recent efforts have adapted VFSS for use in rodents (Lever *et al*., 2015; Mueller et al., 2022; Welby et al., 2020). Here, we used VFSS to analyze mice in which we eliminated selected subtypes of vagal sensory neurons, the ablation of which revealed marked esophageal dysmotility. Our histological and retrograde tracing studies assign this function to the Prox2+/Runx3+ neuronal subtypes MM2 and MM8 that form IGLEs along the entire rostro-caudal axis of the esophagus. Further, the meta-analyses reported here provides information on genes expressed in these neuronal subtypes, and can be mined for the identification of potential drug targets. This might be useful to specifically modulate the vagal sensory neurons that control esophageal peristalsis with the goal to ameliorate motility disorders.

In addition to the vagal sensory arm providing feedback information for esophageal peristalsis, the sensorimotor circuit controlling esophageal motility also includes the hindbrain central pattern generator in the NTS that controls swallowing (Goyal and Chaudhury, 2008; Jean, 2001; Kim *et al*., 2022; Spencer and Hu, 2020). The Prox2+/Runx3+ sensory vagal neurons that we characterized here form synapses with defined neuronal types in the NTS, and in particular we identified Pou3f1+ neurons in the central nucleus of the NTS as synaptic targets. Esophageal afferents are known to project to neurons of previously uncharacterized identity in the central NTS, which in turn project to esophageal motor neurons in the nucleus ambiguus (Altschuler et al., 1989). We suggest that the Pou3f1+ neurons in the central NTS are well positioned to play a key role in esophageal function.

## Acknowledgements

We thank Bettina Brandt and Sven Buchert for technical assistance, and Petra Stallerow and Claudia Päseler for animal husbandry. We also thank the University of Missouri veterinary staff for animal husbandry, and the University of Missouri Pathology laboratory for the monitoring of mice. The authors thank Thomas Müller and Fritz Rathjen for critically reading the manuscript, and the authors of Bai et al., 2019, for providing us with their single cell RNA sequencing data. The authors thank Hans-Peter Rahn of the MDC Flow Cytometry facility, Sandra Raimundo of the MDC Advanced Light Microscopy facility and the staff of the MDC Next Generation Sequencing facility. We thank Mohammed Khallaf and Julia Ojeda-Alonso for help in establishing the *ex vivo* esophagus/stomach vagal nerve preparation. E.D.L. was supported by a Travelling Fellowship from the Company of Biologists (grant number: DEVTF2103518), and an MDC Internal PhD Fellowship. This work was also supported by the Deutsche Forschungsgemeinschaft (DFG, German Research Foundation) under Germanýs Excellence Strategy – EXC-2049 – 39068808 and by the Helmholtz Association (to C.B.).

## Author contributions

Conceptualization: E.D.L. and C.B.; Methodology: E.D.L., P.L.R., A.M., K.S., N.V., K.L.O., X.L., G.R.L., T.E.L.; Software: A.M.; Validation: R.T., T.E.L; Formal Analysis: E.D.L., P.L.R., A.M., S.D.; Investigation: E.D.L., P.L.R., A.M., K.L.O., T.E.L., H.L.; Resources: C.B., J.F.B., A.W., N.Z., R.K., S.J., G.R.L., N.R., T.E.L.; Data Curation: A.M.; Writing – Original Draft: E.D.L. and C.B.; Writing – Review & Editing: E.D.L. and C.B.; Supervision: C.B., S.J., N.R., T.E.L.; Funding Acquisition: C.B.

## Resource Availability

### Lead Contact

Further information and requests for resources and reagents should be directed to and will be fulfilled by the Lead Contact, Carmen Birchmeier.

## Material availability

All unique reagents generated in this study are available from the lead contact with a completed materials transfer agreement.

## Data and code availability

Original datasets supporting the current study are available from the Lead Contact upon request.

## Methods

### Mouse lines

All experiments were conducted according to regulations established by the Max Delbrück Centre for Molecular Medicine, LAGeSo (Landesamt für Gesundheit und Soziales), and the institutional animal care and use committee at the University of Missouri. *Ai65* (#021875), *Ai80* (#025109), *VGlut2^Cre^* (#016963) mice were obtained from the Jackson Laboratory (Daigle *et al*., 2018; Madisen et al., 2015; Vong et al., 2011). *Phox2b^Cre^* (D’Autreaux *et al*., 2011), and *R26^FTLG^* (Dempsey *et al*., 2021) mice were provided by Jean-François Brunet (Institut de Biologie de l’ENS, Paris, France). The *Tau^ds-DTR^* (Britz *et al*., 2015) mice were a kind gift from Martyn Goulding (Salk Institute). *Runx3^Cre^* mice were a kind gift from Yoram Groner (Weizmann Insitute, Rehovot, Israel) (Levanon et al., 2011). The *Gt(ROSA)26Sor<tm2.1Sia>* mice were provided by Shinichi Aizawa (RIKEN Center for Developmental Biology); we refer to them as *R26^nGFP^* mice, as they express a nuclear GFP upon cre-mediated stop cassette excision (Abe et al., 2011). Mice were housed at room temperature (23°C), humidity (56%), and with a 12-hour light-dark cycle.

### Generation of *Prox2^FlpO^* mice

*Prox2^FlpO^* mice were generated at the transgenic core facility at the Max Delbrück Center for Molecular Medicine in Berlin using CRISPR/Cas9 to insert a 1.5kb fragment containing the codon-optimized FLP recombinase (FlpO) and a bovine poly(A) sequence between the 5’ UTR and 1^st^ exon of the *Prox2* gene by homology-directed repair (Wefers et al., 2017). The donor vector was synthesized by GeneArt (Invitrogen GeneArt, ThermoFisher), and contained genomic sequences 2kb upstream (5’ homology arm) and 2kb downstream (3’ homology arm) of the 1^st^ exon of the *Prox2* gene. The successful insertion of the vector was confirmed by long range PCR. *Prox2^FlpO^* mice were born at the correct Mendelian ratios, and could not be distinguished from wild-type littermates based on their appearance, behavior, fertility or lifespan. See also (Nishijima and Ohtoshi, 2006).

### Tissue preparation

Mice were sacrificed and perfused with PBS before organ harvesting. Dissected organs were washed in PBS before fixation in 4% PFA in PBS (1 hour for vagal ganglia, 4 hours for digestive organs and 6 hours for brains). After fixation, organs were washed in PBS, cryopreserved in 15% sucrose overnight at 4°C, and then in 30% sucrose overnight at 4°C. Organs were embedded with Tissue-Tek O.C.T Compound (Sakura) and stored at −80C° until cryosectioning. Vagal ganglia were cryosectioned at 16µm, whereas digestive organs and brains were cryosectioned at 30µm. Sections were stored at −80°C until used.

### Immunohistology

Immunohistology was performed as described with minor modifications (Hernandez-Miranda et al., 2017). In short, sections were thawed, briefly washed (PBS with 0.2% Triton X-100), and blocked (PBS with 0.2% Triton X-100 and 5% normal horse serum) for 1 hour at room temperature. The primary antibody was diluted in blocking solution and incubated for 1-2 days at room temperature. Sections were washed in PBS before being incubated with the secondary antibody diluted in blocking solution for 1 hour at room temperature. Sections were again washed in PBS and mounted with Immu-Mount (ThermoFisher). The following primary antibodies were used: goat anti-Phox2b (R&D Systems, AF4940, 1:200), rabbit anti-RFP (Rockland, 600-401-379-RTU, 1:500), chicken anti-GFP (Aves Labs, GFP-1020, 1:500), rat anti-GFP (Nacalai Tesque, GF090R, 1:1000), goat anti-CD117 (R&D Systems, AF1356, 1:400), sheep anti-nNos (Millipore, AB1529, 1:250), rabbit anti-Tubulin β-3 (BioLegend, Poly18020, 1:1000), goat anti-CTb (List Labs, 703, 1:2000), goat anti-ChAT (Millipore, AB144P, 1:200), rabbit anti-Pou3f1 (Abcam, ab126746, 1:500), and sheep anti-TH (Millipore, AB1542, 1:1000). We used species specific secondary antibodies coupled to Cy2-, Cy3-and Cy5 (Jackson ImmunoResearch, 1:500).

### Single molecule fluorescent *in situ* hybridization (RNAscope)

*In situ* fluorescent hybridization was performed using the RNAscope Multiplex Fluorescent Reagent Kit V2 from ACDbio according to the manufacturer’s instructions. Briefly, vagal ganglia sections were thawed at 37°C for 20 minutes and post-fixed in 4% PFA in PBS for 15 minutes before washing in PBS and continuing with the manufacturer’s instructions. For combining immunohistology with RNAscope we proceeded until the hydrogen peroxide wash, then washed the sections in PBS and incubated them at 4°C overnight with the primary antibody diluted in Co-Detection Antibody Diluent (obtained from ACDBio). The sections were washed in PBS and the protease treatment was performed using Protease III. The RNAscope protocol was then continued, the sections washed in PBS and incubated with the secondary antibody for 1 hour at room temperature in blocking solution (PBS with 0.2% Triton X-100 and 5% normal horse serum). Sections were washed in PBS and mounted with ProLong Gold Antifade mountant (ThermoFisher). We used the following probes in this study: Prox2 (593331-C3), Runx3 (451271 and 451271-C2), Piezo2 (400191-C2 and 400191-C3), Trpv1 (313331), Prrxl1 (446631), Calb1 (428431-C3), Adra2a (425341-C3), Slc18a3 (448771-C3), Slc17a6 (319171-C3), Phox2b (407861-C2 and 407861-C3), Rbp4 (508501-C2), Gata3 (403321), Grm5 (423631-C2), Lamp5 (451071-C2), Mc4r (319181), Gabrg1 (501401-C3), and Glp1r (418851).

### Vagal neuron isolation

We dissected the vagal ganglia from 15 *VGlut2^Cre^;R26^nGFP^* mice of either sex at P4, removed excess nerve, muscle and vascular tissue, and placed them in a 1.5 ml Eppendorf tube with warm F12/FHS (F12 with 10% fetal horse serum). Neurons were isolated essentially as described (Lechner and Lewin, 2009). In short, ganglia were digested in F12/FHS solution containing 0.125% collagenase, incubated at 37°C for 1 hour, washed 3x in PBS, and then incubated in PBS with 0.25% trypsin at 37°C for 15 minutes. Ganglia were dissociated using fire-polished Pasteur pipettes of decreasing diameter. The solution was transferred on top of a 2 ml BSA cushion (F12/FHS solution with 15% bovine serum albumin) and spun for 10 minutes at 900 RPM. The cell pellet was resuspended in 500 µl of HBSS without calcium or magnesium, strained twice through a 70 µm filter (Sysmex), and DAPI (Sigma) was added to a final concentration of 300 nM to label dead cells before sorting. We sorted GFP-positive / DAPI-negative cells into 96-well plates using ARIA Sorter III (BD) and BD FACSDiva software 8.0.1.

### Library generation and sequencing

Single cell RNA sequencing was done following the CEL-Seq2 protocol (Hashimshony *et al*., 2016). cDNA Libraries were prepared for 16 96-well plates, pooled together, and sequenced on a NextSeq 500 (Ilumina) in two separate runs by the next generation sequencing core facility of the Max-Delbrück Center for Molecular Medicine.

### Clearing and whole organ immunohistology

Mice were sacrificed, perfused with PBS to remove the blood and then perfused with 4% PFA in PBS. Vagal ganglia and digestive organs were dissected and fixed overnight at 4°C in 4% PFA in PBS. Tissue was cleared using the CUBIC protocol (Susaki *et al*., 2015). Briefly, tissue was first washed overnight at room temperature in PBS and then immersed in Sca*l*eCUBIC-1 (a mixture of 25% urea, 25% Quadrol, 15% Triton X-100 and 35% MQ-H_2_O) diluted 1:1 with MQ-H_2_O overnight in a 37°C water bath. The tissue was then placed in Sca*l*eCUBIC-1 in a 37°C water bath until the tissue became transparent (1/2 a day for vagal ganglia and 2-4 days for digestive organs). The solution was exchanged every 2 days. Once the organs were sufficiently cleared they were washed overnight at room temperature in PBS + 0.2% Triton X-100. Next, we incubated the organs with the primary antibodies diluted in modified blocking solution (PBS with 10% Triton X-100, 5% normal horse serum and 300mM NaCl) for 10 days on a shaker at 37°C, refreshing the antibodies after 5 days. Organs were washed overnight in PBS with 0.2% Triton X-100, incubated with the secondary antibodies and DAPI in modified blocking solution for 8 days on a shaker at 37°C, refreshing the antibodies after 4 days. The tissue was then immersed in EasyIndex RI = 1.46 (LifeCanvas Technologies) diluted 1:1 with MQ-H_2_O overnight in a 37°C water bath, before being transferred to EasyIndex RI = 1.46 overnight for the final refraction index matching. Once the tissue was cleared, stained and refractive index matched it was placed into a square plastic mold filled with a 2% low melting point agarose prepared with EasyIndex RI = 1.46.

Cleared ganglia were imaged using a Zeiss lightsheet 7 microscope, while cleared digestive organs were imaged using a custom built mesoSPIM microscope (Voigt *et al*., 2019). Cleared, stained and embedded digestive organs were immersed in EasyIndex RI = 1.46 inside a small quartz glass cuvette (45 x 12.5 x 22.5 mm Portmann Instruments AG, UQ-205, quartz glass), which was placed inside a larger chamber (40 x 40 x 100 mm, Portmann Instruments AG, UQ-753-H100) filled with RI-matching liquid for fused silica (RI=1.46, Cargille Cat. #19569). The lightsheet illumination was delivered sequentially from the left and right sides, resulting in two different stacks per view, which were registered and fused into a single stack using BigStitcher (Horl et al., 2019) for each channel. The excitation laser line (Hübner Photonics C-Flex: 561 nm) was used with the corresponding detection filter (Chroma ET590/50m). The esophagus/stomach from an adult (3 months of age) *Prox2^FlpO^;Phox2b^Cre^;Ai65* (*Prox2/Runx3^Tom^*) mouse was acquired with a 1 x 3 tile scan at 1x zoom, while the esophagus/stomach from an adult (3 months of age) *Runx3^Cre^;Prox2^FlpO^;Ai65* (*Runx3^Tom^*) mouse was acquired with a 2 x 6 tile scan at 2x zoom (Olympus 1x MVPLAPO1x). The images displayed in Figure 2A are maximum intensity Z-projections of the final fused images. Vagal neuron counts at P4 were performed with Imaris v9.9.

### Esophagus and stomach single fiber nerve recordings and analysis

Electrophysiological single fiber nerve recordings were realized using an *ex-vivo* esophagus/stomach vagal nerve preparation (Page and Blackshaw, 1998). Briefly, adult *Prox2/Runx3^ChR^* (*Prox2^FlpO^;Phox2b^Cre^;Ai80*) mice were sacrificed by CO_2_ euthanasia. Mice were transcardially perfused (40 ml) with a carbogen equilibrated extracellular solution (125 mM NaCl, 2.5 mM KCl, 25 mM NaHCO_3_, 1.25 mM NaH_2_PO_4_, 1 mM MgCl_2_, 2 mM CaCl_2_, 20 mM glucose and 20 mM HEPES). The entire esophagus or stomach was pinned in a dissection chamber with flowing equilibrated extracellular solution (20-23°C), with the attached vagus nerve that was isolated from the cervical to the thoracic esophagus (esophagus analysis) or from the cervical esophagus to the lower esophageal sphincter (stomach analysis). For the esophagus analysis, an open-book preparation was obtained by sectioning the esophagus longitudinally, taking care not to damage lateral ramifications of the vagus nerve. Esophageal recordings were made by probing the thoracic to abdominal esophagus. For the stomach preparation, the esophagus was removed at the lower esophageal sphincter, and the intestine was removed 1.5-2cm distal to the pyloric sphincter. A cannula (0.63 mm diameter) connected to a 1 ml syringe filled with fresh extracellular solution was inserted into the intestine and secured in place with a suture. A small hole was perforated in the forestomach with a 0.8 mm gauge needle to allow for pressure equilibration after stomach distension. The preparation was then transferred to a recording chamber perfused with warm (32°C) carbogen saturated extracellular solution, and the vagus nerve was passed through a channel to an adjacent recording chamber containing mineral oil. Small nerve bundles were teased apart and placed on a platinum recording electrode.

In the esophagus and stomach, mechanosensitive fibers were identified by manually poking the tissue in the recording chamber, and tested for their Prox2/Runx3 identity using optogenetic stimulation (470 nm Thorlab diode laser, 1 second stimulation, intensity 2 mW/mm^2^) delivered by a 1 mm fiber optic placed on the responsive field. After observing a light-sensitive response, mechanosensitive esophageal fiber characterization was performed using a piezo actuator (Physik Instrumente, Germany, P-602.508) connected to a force measurement device (Kleindiek Nanotechnik, Reutlingen, Germany, PL-FMS-LS) (Walcher et al., 2018). Different mechanical forces (ranging from 50 mN to 250 mN) were applied with a 2 sec static phase. We allowed fibers to recover for 1 min before applying the next stimulation. For analysis of stomach mechanosensitive fibers, we applied increasing volumes (0.1, 0.2, 0.3 ml) to distend the entire stomach, allowing pressure equilibration by removing the added liquid and waiting 5 minutes between each distension. Stomach distention was continuously recorded via a force measurement device connected to a 1 mm probe placed on the ventral wall of the stomach.

Raw data were recorded using an analog output from a Neurolog amplifier, filtered (100-5 kHz) and digitized (10 kHz) using a Powerlab 4/30 system and Labchart 8 software with the spike-histogram extension (ADInstruments Ltd., Dunedin, New Zealand). Single nerve fiber recordings were further analyzed to identify single units using Spike2 software (ced.co, Spikes2 version 10). Individual spike units were identified based on the following parameters: new template width as % of amplitude: 32, minimum % of points in template: 60, minimum occurrence of events: 1/50, spike duration considered for the sorting −1.5 to 1.5 ms. Unit spikes were binned per 200 ms to ease data analysis and light/mechanosensitive and -insensitive units were separated. For each fiber recording, between 3 - 10 separate mechanosensitive units were identified. To be categorized as Prox2/Runx3-positive, the unit had to increase its firing at least 2-fold during light stimulation. For each preparation, between 4-7 light stimulations were performed. To be considered Prox2/Runx3-positive, units had to respond to at least 60% of the stimuli.

Firing patterns were further analyzed using Microsoft Excel. Prox2/Runx3-positive and -negative units were aligned using the onset of the mechanical stimuli, and activities of individual units were normalized using the basic activity measure of the unit 2 seconds before the stimulus onset to obtain the Δ Frequency (Hz). For population quantifications, mechanosensitive, light sensitive, and mecho- and light sensitive units were binned, and the unit activity observed 1 second after the onset of the stimuli was averaged. Prox2/Runx3-positive fibers were manually decomposed depending on their firing pattern during either the 300 mN mechanical or the 0.3 ml stomach distention. Two types of firing responses were observed, which were separately binned using a firing decay of 40% 1 second after the stimulus onset as criterion. Type I units displayed a prolonged firing pattern with a slower decay, i.e. the units continued to spike until the offset of the mechanical stimuli (slowly adapting). Type II units displayed a fast decay after the stimulus onset, and rapidly returned to basal activity before the end of the stimuli (rapidly adapting).

### Retrograde tracing

#### CTb tracing from the abdominal esophagus and stomach

WT adult mice (∼3 month of age) were anesthetized with an intraperitoneal injection of ketamine/xylazine (80 mg/kg body weight ketamine and 10 mg/kg body weight xylazine). The abdomen was shaved and betadine was applied to the abdominal skin. Ophthalmic ointment was applied to the eyes to prevent them from drying out during the procedure. A small transverse laparotomy was made below the sternum and fire polished glass Pasteur pipettes were used to position the organs prior to injection. Glass injection needles were prepared with a DMZ universal electrode puller (Zeitz-Instruments) and were filled with 0.5% CTb solution (ThermoFisher) in 0.9% NaCl. We added 0.2% Fast Green (Sigma) in order to visualize the injection site and confirm a successful injection. For glandular and non-glandular stomach injections a total of 2 µl was injected, while for abdominal esophagus 1 µl was injected, due to the smaller size of the target region. The injections were performed using a Nanoject III Programmable Nanoliter Injector (Drummond Scientific Company), with a volume of 250 nl and a speed of 50 nl/s per injection. In all cases, the needle was carefully placed into the muscle layer of the target regions and allowed to remain in place for 10 s before and after each injection. After the injections we performed layered wound closure, suturing the abdominal muscle layer first, followed by suturing the skin. Post-surgery mice were given a subcutaneous injection of Carprofen (5 mg/kg body weight) and received Metamizol 1 ml/100 ml drinking water for analgesia. Mice were monitored twice daily to ensure that they were recovering properly, then were euthanized 4-5 days following surgery for vagal ganglia harvesting.

#### Fast Blue tracing from the cervical and thoracic esophagus

We first measured the length of the esophagus in P7 WT mice and found that it measured around 2 cm from the base of the tongue to the stomach. Next, we prepared a 0.25% Fast Blue solution in 50% 0.9% NaCl and 50% glycerol. To trace from the cervical and thoracic esophagus, mice were gavaged 2 times with 2 µl of the Fast Blue solution 0.5 cm and 1.3 cm from the base of the tongue, respectively. Animals were monitored twice daily and were euthanized 7 days following Fast Blue gavage.

### Analysis of the P4 vagal ganglia CEL-Seq2 data

Data processing and gene quantification was performed using dropseq-tools v2.0 and picard-tools v2.18.17. We used the standard pipeline for Drop-seq data, with the necessary adaptations to analyze the CEL-Seq2 data. We removed the bead barcode correction steps, and in the DigitalExpression quantification we inputted the list of 96 CEL-Seq2 barcodes. Alignment was performed with STAR v2.5.3a. We used the GRCm38 genome and the annotation from the GRCm38.p4 assembly. Two thresholds were set to filter out wells without cells or wells with multiple cells. We set a lower threshold of 17,000 UMIs (unique molecular identifier) per cell, and an upper threshold of 250,000 UMIs per cell. These UMI thresholds filtered out 144 cells, leaving 1392 out of 1536 cells (from 16 96-well plates). We removed 138+6=144 cells (9.4% of the total cell count).

After quantification, downstream analysis was performed with Seurat v3.0. Seurat was run with the default parameters with the following exceptions. 2000 genes were selected with the FindVariableFeatures function of Seurat. The first 30 principal components were selected after PCA, excluding PC3 and PC11 as these PCs represented glial contamination in our dataset. The neighbor graph was constructed with FindNeighbors with a k parameter of 11. The clustering resolution was set to 1. UMAP visualization was used with the correlation metric and 20 number of neighbors.

### Integration analysis

For the integration analysis the raw count tables were used from the Ernfors and Knight laboratories (Bai et al., 2019; Kupari et al., 2019). The first was downloaded from a public repository and the second was kindly provided by the authors after our request. The Bai et al., 2019 data consisted of two datasets, the targeted and the untargeted cells, which were sequenced with different protocols (Bai et al., 2019). Seurat v3.0 canonical correlation was used to integrate the four datasets (including ours), using 30 canonical components. Initially, PCA was performed for 30 principal components. The neighbor graph was constructed with those 30 components, with a k parameter of 20, and clustering was performed with a resolution of 1. After this initial analysis we identified clusters of glial and epithelial cells and sympathetic neurons. We then removed these cell clusters and repeated the integration analysis on the remaining cells. This time integration was performed using 40 canonical components. After PCA, the first 29 components were used, excluding principal components 4 and 6 as these carried a glial signal. The rest of the downstream integration analysis was performed using default parameters.

### Ablation

We generated *Prox2/Runx3^ds-DTR^*(*Prox2^FlpO^;Phox2b^Cre^;Tau^ds-DTR^*) mice in which all cells with a history of Phox2b and Prox2 expression expressed the human diphtheria toxin receptor. We ablated *Prox2+*/*Runx3+* vagal neurons in adult mice (∼6 months of age) by i.p. injection of Diphtheria toxin (DT, Sigma), reconstituted in 0.9% NaCl, and injected at a concentration of 40 ng per gram bodyweight. After DT administration the ablation mice were monitored twice daily. If their weight dropped below 10% of their starting weight, they received a 0.5 ml i.p. injection of 0.9% NaCl daily and were given access to a nutritionally fortified water gel (DietGel Recovery from ClearH_2_O). One male mouse continued to lose weight after the ablation and was euthanized before it could undergo endline testing. All other mice stabilized between 5-7 days post ablation.

### Behavior

Videofluoroscopy experiments to measure *in vivo* swallowing function were performed as previously described (Lever *et al*., 2015; Welby *et al*., 2020). Briefly, mice (n=15, 8M and 7F, genotype: *Prox2^FlpO^;Phox2b^Cre^;Tau^ds-DTR^*, 6 months of age) underwent videofluoroscopy swallow study (VFSS) at the University of Missouri using customized equipment and analysis software. The mice were shipped from the Max Delbrück Center in Berlin, Germany to the University of Missouri in MO, USA and were placed in quarantine for 3 weeks. During this time the mice were behaviorally conditioned with the VFSS chamber and oral contrast solution in order to familiarize them with the experimental set up and facilitate drinking during VFSS testing. Following release from quarantine, the mice were water restricted overnight for 12 hours in order to increase their motivation to drink during testing the following morning. During the water restriction period, a VFSS chamber was placed in each home cage. Mice were each tested in their respective home cage chambers the following morning. VFSS testing was performed individually using a miniaturized, low energy (30 kV, 0.2 mA) fluoroscope (The LabScope, Glenbrook Technologies, Newark, NJ, USA) and videos were captured at 30 frames per second (fps). Mice were gently placed into the VFSS chamber and enclosed using two end-caps. One end-cap had a small bowl attached through which the liquid contrast agent (Omnipaque, GE Healthcare, 350 mg iodine/mL; diluted to a 25% solution with deionized water and 3% chocolate flavoring) could be administered during testing. The test chamber was then positioned within the lateral plane of the fluoroscope, and the bowl was filled with the liquid contrast agent via a custom syringe delivery device. When the mice began to drink, the fluoroscope was activated via a foot pedal. In order to minimize the radiation exposure time, the fluoroscope was turned off when the mice turned away from the bowl or initiated non-drinking behaviors. If mice did not drink, we placed them back in their home cage for ∼30 mins and re-tested them. We captured drinking bouts in videos of approximately 30-60 s duration and saved them as AVI files. After baseline behavioral testing, mice underwent DT ablation of *Prox2+* and *Runx3+* vagal neurons. We allowed the mice to recover for 19 days after the ablation, at which point we re-tested the mice using the same VFSS protocol as above. Thus, we could compare the swallowing behavior of the same mice at baseline (pre-DT ablation) versus endline (post-DT ablation).

We imported the AVI files into Pinnacle Studio (version 24; Pinnacle Systems, Inc., Mountain View, CA) and identified drinking bouts between 2-5 s in length to obtain a total of 3-15 s of uninterrupted drinking per mouse. The start of the drinking bouts was always at a swallow event, defined as when the liquid bolus abruptly moved from the vallecular space to the esophagus. The 2-5 s video clips were then imported into a custom VFSS analysis software (JawTrack™, University of Missouri) for subsequent frame by frame analysis. The software allows for the semi-automated extraction of many swallowing parameters such as lick rate (number of jaw open/close cycles per second, calculated for 2-5 s long video clips and then averaged), lick interval (time between successive lick cycles throughout a 2-5 s long video clip, and then averaged), swallow rate (number of swallows in each second of a 2-5 s video clip, then averaged), swallow interval (time between successive swallows throughout a 2-5 s video clip, then averaged), lick-swallow ratio (number of jaw open/close cycles between each successive swallow pair throughout a 2-5 s video clip, then averaged), pharyngeal transit time (bolus travel time through the pharynx for each swallow, then averaged), esophageal transit time (bolus travel time through the esophagus, then averaged), jaw closing velocity (speed at which the jaw closes during each jaw cycle throughout a 2-5 s video clip, then averaged) and jaw opening velocity (speed at which the jaw opens during each jaw cycle throughout a 2-5 s video clip, then averaged) (Mueller *et al*., 2022; Welby *et al*., 2020).

### Quantification and statistical analyses

All details regarding the number of samples and statistical tests performed can be found in the figure legends. All paired and unpaired t-tests were two-tailed, and no statistical test was done to predetermine sample sizes. One animal (#5697) was removed as an outlier from the Swallow interval and Lick to swallow ratio plots (Figure S5C), as it failed the Grubbs’ test with alpha = 0.01. Significance for t-tests and ANOVAs was defined as * p < 0.05, ** p < 0.01, *** p < 0.001, **** p < 0.0001. Statistical analyses were performed using GraphPad Prism v6.0c.

---

**Figure S1. scRNAseq analysis assigns a developmental origin and putative function to vagal neuron subtypes, related to** Figure 1.

**(A)** Lightsheet imaging (acquired with a Zeiss lightsheet 7) of a vagal ganglion from a *VGlut2^Cre^;R26^nGFP^* mouse at postnatal day P4; GFP+ neuronal nuclei (green, **top**), and all detected nuclei (gray, **bottom**) are shown. Detected nuclei were quantified using IMARIS v9.9.0 (5288±423 nuclei per ganglion, n = 3). **(B)** UMAP plot based on single cell RNA-seq data from 1392 neurons isolated at P4 from *VGlut2^Cre^;R26^nGFP^* animals revealed 22 clusters (color coded and numbered). **(C)** UMAP plot based on our meta-analysis of 4,442 neurons revealed 27 clusters (color coded and numbered, *M*eta-*J*ugular (MJ) 1-6, *M*eta-*C*hemo (MC) 1-11, *M*eta-*M*echano (MM) 1-10). **(D)** UMAP plot based on the meta-analysis displaying cells of each data set by separate colors; all data sets contributed cells to each cluster. **(E)** smFISH showing that all vagal neurons are *VGlut2*+ (Slc17a6); they differ in *Prrxl1* and *Phox2b* expression that differentiates jugular and nodose ganglia, respectively. **(F)** smFISH showing that all nodose neurons are *Phox2b*+; they differ in *Piezo2* and *Trpv1* expression that mark putative mechanosensory and chemosensory neurons, respectively. **(G)** UMAP plots for *VGlut2* (*Slc17a6*), *Prrxl1*, *Phox2b*, *Trpv1* and *Piezo2*. Data in A are represented as mean ± SD.

**Figure S2. Generation and characterization of the *Prox2^FlpO^* mouse strain, related to** Figure 2**. (A)** Scheme showing the strategy used to generate the *Prox2^FlpO^* allele. **(B)** *In situ* hybridizations against *Prox2* mRNA in sagittal sections of E11.5 (left) and E13.5 (right) embryos; pictures were obtained from the Allen Brain Atlas. **(C)** Recombination efficiency of *Prox2*+ neurons in *Prox2/Runx3^Tom^* (*Prox2^FlpO^;Phox2b^Cre^;Ai65*) **(left)** and *Runx3^Tom^* (*Runx3^Cre^;Prox2^FlpO^;Ai65*) **(right)** animals. Quantifications show that 94.8±3.6% and 11.3±3.6% (p < 0.0001) of *Prox2*+ neurons were recombined in *Prox2/Runx3^Tom^* and *Runx3^Tom^* mice, respectively, n = 4. **(D)** Recombination efficiency in *Runx3*+ neurons in *Prox2/Runx3^Tom^* **(left)** and *Runx3^Tom^* **(right)** animals, n = 4. Quantifications show that 71.4±2.1% and 70.5±4.8% (p = 0.744) of *Runx3*+ neurons were recombined in *Prox2/Runx3^Tom^* and *Runx3^Tom^* mice, respectively, n = 4. **(E)** Representative smFISH images from *Prox2/Runx3^Tom^***(left)** animals showing recombination specificity in Prox2+ (green) and Runx3+ (red) neuronal subtypes and *Runx3^Tom^* **(right)** animals showing recombination specificity in Runx3+ (red) subtypes, quantified below, n = 4. **(F)** Proposed lineage tree of Prox2 and Runx3 neurogenesis. Phox2b+ nodose neurons are born during early development and first express *Prox2* before largely separating into Prox2+ and Runx3+ subtypes. Thus, Runx3+ neurons have a history of Prox2 expression and can be labelled by the used intersectional strategy (*Runx3^Cre^;Prox2^FlpO^*). **(G)** Labelling strategy to mark Prox2+ and Runx3+ vagal neurons with Tomato using *Prox2/Runx3^Tom^* (*Prox2^FlpO^;Phox2b^Cre^;Ai65*) mice. **(H)** Labelling strategy to mark Runx3 vagal neurons with Tomato using *Runx3^Tom^* (*Runx3^Cre^;Prox2^FlpO^;Ai65*) mice. Data are represented as mean ± SD, ****p < 0.0001, unpaired two-tailed t-test.

**Figure S3. MM1 and MM8 are *Glp1r*+ stomach projecting neurons, related to** Figure 3.

**(A)** smFISH **(left)** analysis using a *Piezo2* (red) probe, combined with an immunohistological analysis of CTb (green) after CTb injection into the glandular stomach. **(Right)** quantification of *Piezo2*+ neurons identified after CTb or Fast Blue injection into various sites of the upper gastrointestinal tract. *Piezo2*+ neurons corresponded to 53.4±19.2%, 56.1±20.2%, 17.3±9.5%, 26.8±6.7%, and 31.3±4.3% of all neurons traced from the cervical, thoracic, abdominal esophagus, and non-glandular and glandular stomach, respectively, n = 3 – 4. Note that neurons from the cervical and thoracic esophagus were traced using Fast Blue at P7, and neurons from the abdominal esophagus, non-glandular and glandular stomach were traced using CTb in adults. **(B)** smFISH analysis **(left)** of *Piezo2* (green) and *Glp1r* (blue) mRNA, combined with immunohistological analysis of CTb (red) after CTb injection into the glandular and non-glandular stomach; the quantification is shown on the **right**. *Glp1r*+ neurons corresponded to 77.3±7.4% and 87.7±11.0% of all *Piezo2*+CTb+ neurons traced from the non-glandular and glandular stomach, respectively, n = 4. **(C)** smFISH **(left)** analysis identifying MM1 neurons (probes: *Piezo2* in green, *Rbp4* in red, and *Glp1r* in blue) shows that 89.6±5.3% of MM1 neurons are *Glp1r*+, n = 4. smFISH **(middle)** analysis identifying MM8 neurons (probes: *Adra2a* in green, *Runx3* in red, and *Glp1r* in blue) shows that 94.3±2.6% of MM8 neurons are *Glp1r*+, n = 4. smFISH **(right)** for MM2 neurons (probes: *Piezo2* in green, *Grm5* in red, and *Glp1r* in blue) shows that most MM2 neurons are *Glp1r-*negative: only 2.4±3.0% of MM2 neurons expressed *Glp1r*, n = 4. Data are represented as mean ± SD, ****p < 0.0001, unpaired two-tailed t-test **(B)**, Ordinary one-way ANOVA with Tukey’s multiple comparisons test **(C)**.

**Figure S4. Optogenetic activation of Prox2+/Runx3+ mechanoreceptors, related to** Figure 4**. (A,B)** Quantification of the firing properties of Prox2/Runx3-positive (red, light-sensitive) and Prox2/Runx3-negative (gray, light-insensitive) single fibers. Action potential firing frequency (Hz) was measured on single esophageal **(A)** or stomach **(B)** fibers from *Prox2/Runx3^ChR^* (*Prox2^FlpO^;Phox2b^Cre^;Ai80*) animals with and without blue light stimulation (shown in blue shading). Note that under light stimulation, Prox2/Runx3-positive units dramatically increased their firing frequency (esophagus light sensitive units: 1.4±0.9 Hz to 21.6±3.2 Hz, p < 0.0001, stomach light sensitive units: 2.5±0.7 Hz to 17.6±2.2 Hz, p < 0.0001), n = 6. **(C)** Spike sorting identified three individual mechanosensitive units (shown in red, blue and green) present in a single fiber. Points form a raster plot of single unit instantaneous firing frequencies after stomach distention (**left**, 0.3 ml stimuli, the black line underneath the plot displays the change in stomach volume), or light activation (indicated by blue shading). Note that units 1 and 2 (red and blue) are light-sensitive and thus Prox2/Runx3-positive, whereas the light insensitive unit 3 (green) is Prox2/Runx3-negative. Cycle trigger average (CTA) of 500 spikes (right) of the 3 units identified in this fiber after stomach distension. Data are represented as mean ± SD, ****p < 0.0001, Ordinary one-way ANOVA with Tukey’s multiple comparisons test.

**Figure S5. Oropharyngeal swallow-related behaviors were unaffected after Prox2+/Runx3+ ablation, related to** Figure 5**. (A)** Graph showing weight change in *Control^ds-DTR^* (*Prox2^FlpO^;Tau^ds-DTR^*) control mice for the 14 days following DT administration. **(B)** Outline of the experimental timeline. **(C)** Many oropharyngeal swallow-related behaviors were unaffected after Prox2+/Runx3+ ablation. Swallow rate (2.4±0.4 swallows / s to 2.2±0.5 swallows / s), swallow interval (0.5±0.1 s to 0.5±0.1 s), lick to swallow ratio (3.1±0.9 # of licks / # of swallows to 3.3±1.0 # of licks / # of swallows), jaw opening velocity (20.3±3.4 mm / s to 19.3±4.3 mm / s) and jaw closing velocity (21.3±3.4 mm / s to 19.6±4.8 mm / s) were all unaffected, n = 13 - 14. Data are represented as mean ± SD, paired two-tailed t-test.

