## Supplemental Figures for "Prox2+ and Runx3+ neurons regulate esophageal motility"

### Figure S1 - scRNA-seq analysis assigns a developmental origin and putative function to vagal neuron subtypes

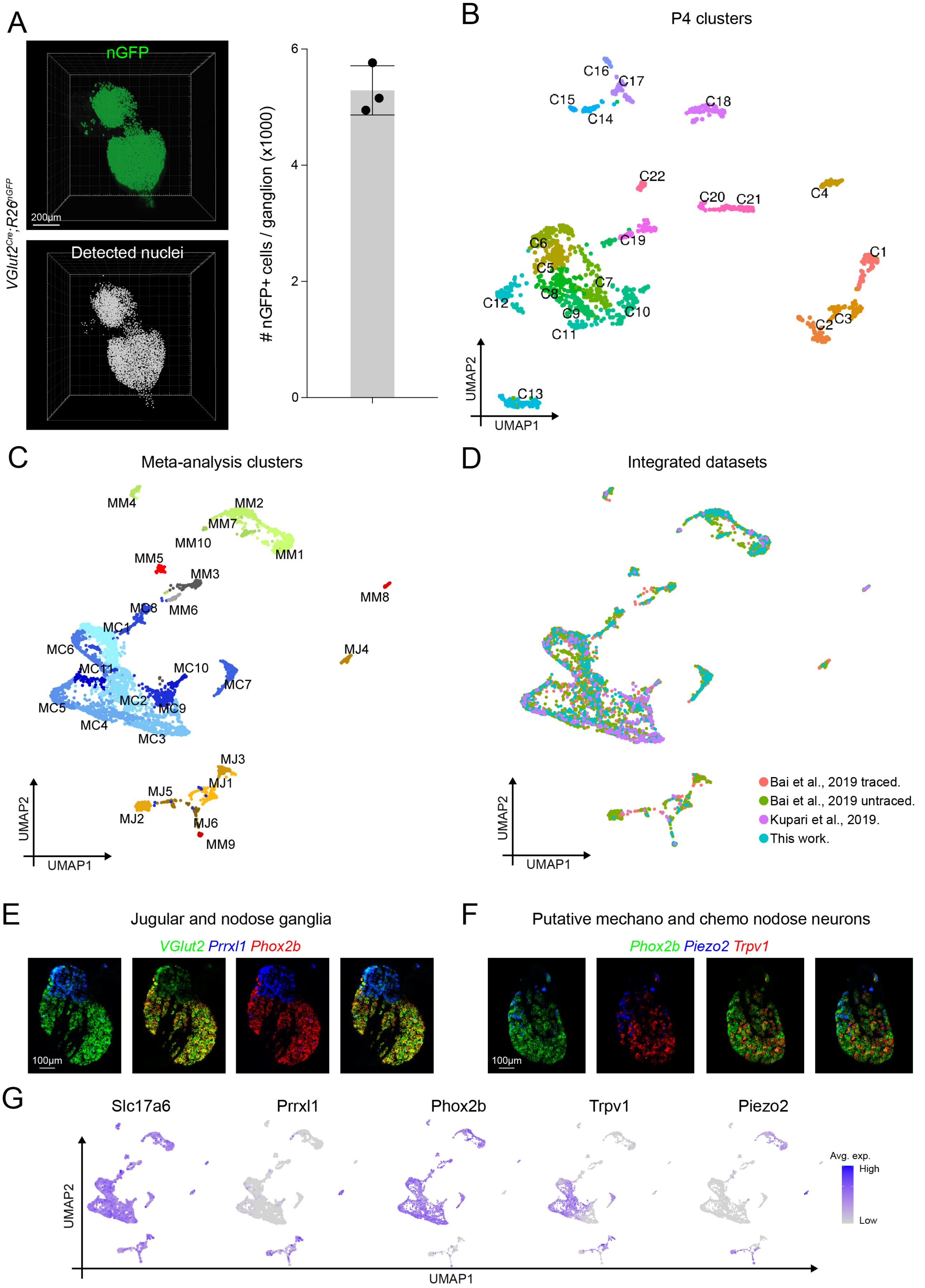

### Figure S2 - Generation and characterization of the *Prox2*$^{FlpO}$ mouse strain

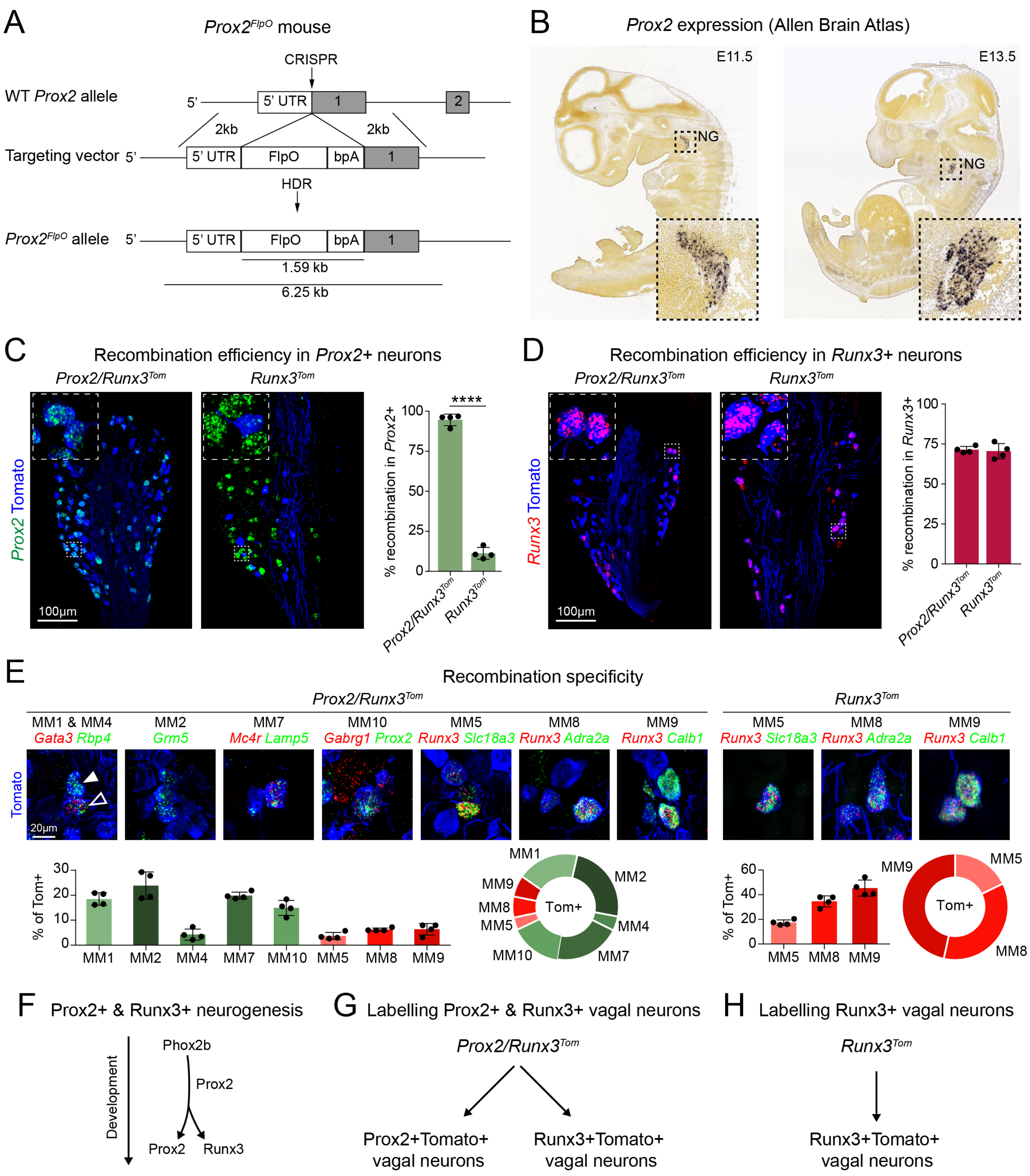

### Figure S3 - MM1 and MM8 are *Glp1r*+ stomach projecting neurons

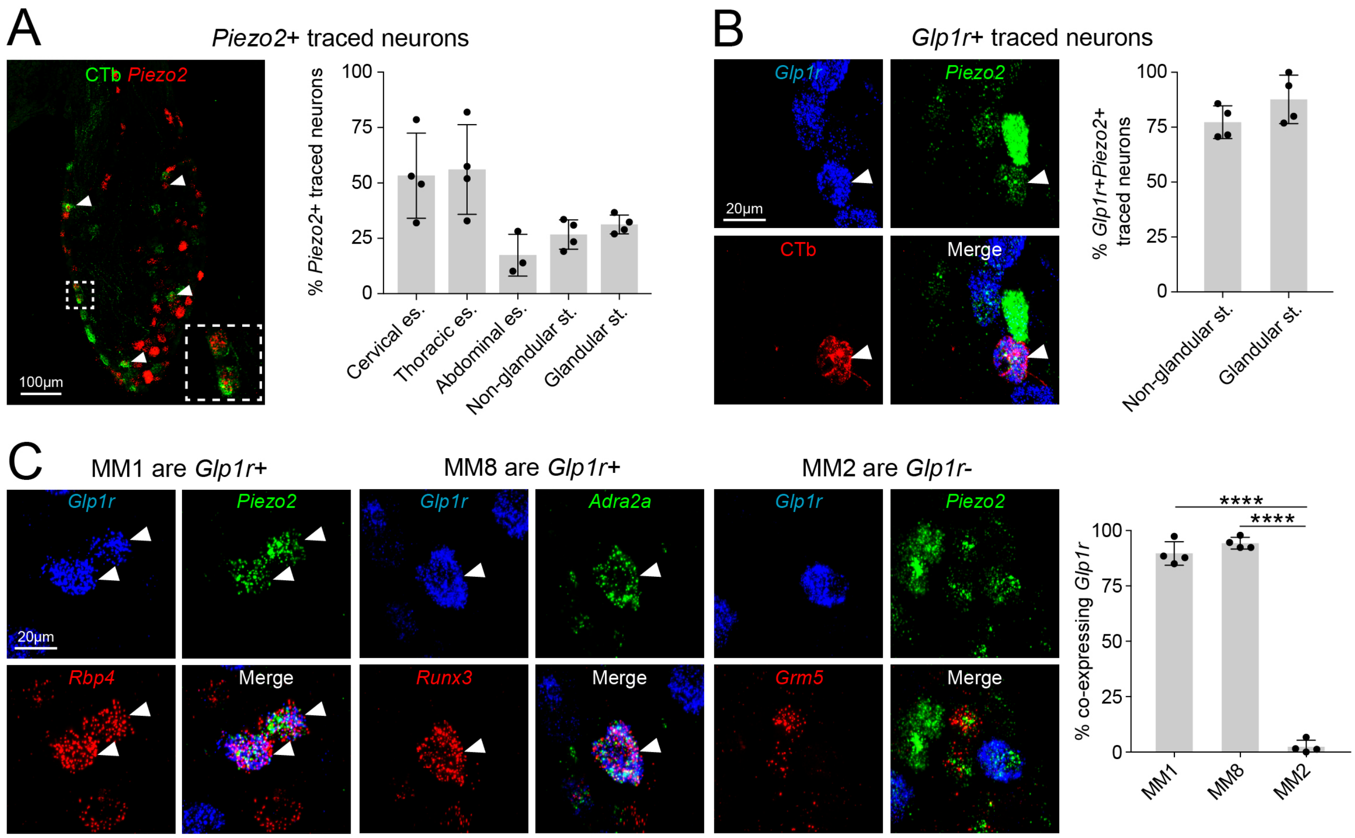

### Figure S4 - Optogenetic activation of Prox2+/Runx3+ mechanoreceptors

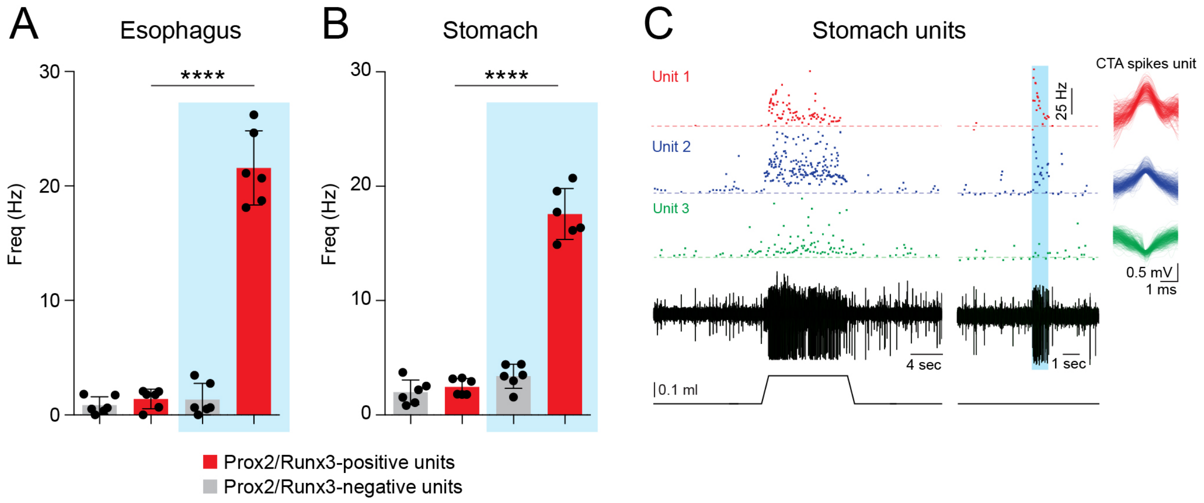

Figure S5 - Oropharyngeal swallow-related behaviors were unaffected after Prox2+/Runx3+ neuronal ablation

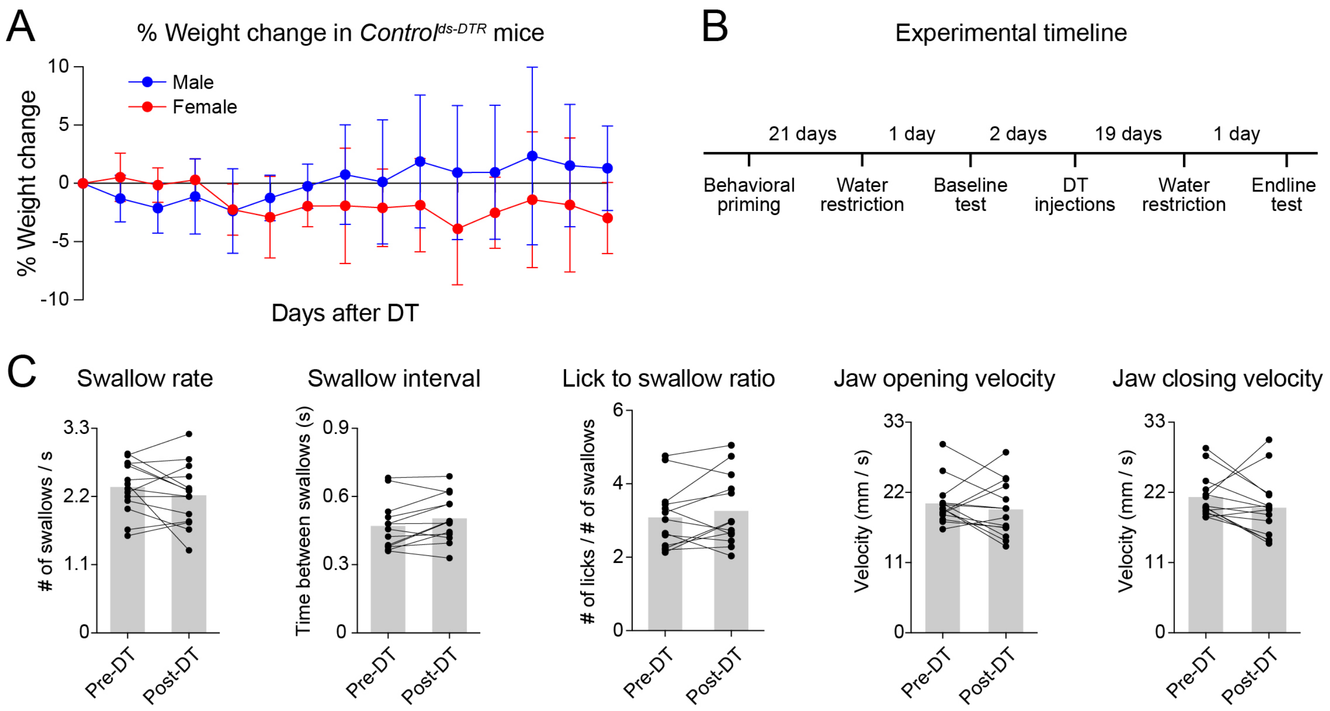
